# Chemical diversity promotes ecosystem function

**DOI:** 10.1101/2025.02.16.638531

**Authors:** Jeremy A. Fonvielle, Sarah R. Sandor, Thorsten Dittmar, Andrew J. Tanentzap

## Abstract

Nature benefits people in many ways that rely on how microorganisms and abiotic processes interact with organic matter to influence ecosystem function (EF). Despite many studies evaluating how biodiversity affects EF, most variation in EF remains unexplained. Here, we tested how the variety of organic compounds in the environment, termed chemodiversity, influences EF. Using a laboratory experiment and 101-lake survey spanning Europe, we discovered that chemodiversity, not biodiversity, consistently explained variation in multiple functions related to carbon cycling and ecosystem metabolism. We attributed positive chemodiversity-ecosystem multifunctionality relationships to the accumulation of compounds that could be biochemically transformed in more diverse ways at higher chemodiversity. We estimated chemodiversity will decline by 2100 due to changes in organic matter sources, causing European lakes to emit, on average, 1.6-times more carbon. These findings reframe our understanding of ecosystem functioning and suggest chemodiversity is key to sustain the benefits that nature delivers to people.

## Introduction

Nature benefits society such as by regulating the climate and providing food and clean water^1,2^. These services are in part delivered by the multiple functions undertaken by microorganisms that break down and transform different organic substrates^3^. For example, as microbes assimilate detritus and increase in biomass^4^, they support the growth of higher trophic levels^5^, as well as modify organic compounds into forms that sequester carbon longer in the environment and thereby contribute to climate regulation^6^. Consequently, ecosystem function is enhanced by both a greater diversity of metabolic capabilities in microbial communities^3^ and the environmental conditions that favour this biodiversity^7^. However, large amounts of variation in the capacity of ecosystems to provide multiple functions simultaneously, hereafter termed ecosystem multifunctionality^1^, remain unexplained. Many large-scale studies have found that biodiversity only explains between 2 to 20% of the variation in ecosystem multifunctionality^3,8,9^, though there are some exceptions^10^.

The organic substrates that microbes use to influence biological, geochemical, and physical processes within ecosystems are an overlooked source of variation in past studies that predict ecosystem multifunctionality^6,11,12^. Natural organic matter in aquatic and terrestrial ecosystems contains tens of thousands (and likely millions) of unique molecules of varying origin and composition, making it one of the most complex mixtures on Earth^13^. This variety of molecules within organic matter and their characteristics, such as their bioavailability and stoichiometry, are together referred to as chemodiversity^14^. Chemodiversity can be described by measuring the diversity and abundance of molecular formulae and their characteristics, similar to approaches for studying biological mixtures of species^15^. Despite many surveys of chemodiversity in waters and soils^16,17^, there has been no attempt to ask how chemodiversity influences ecosystem function in nature as compared with biodiversity and other environmental variables.

We hypothesised that chemodiversity influences ecosystem multifunctionality in two main ways. First, the chemical properties of organic compounds (e.g. structure, elemental composition, and biological and photochemical reactivity) determine the biotic and abiotic pathways by which they are degraded and transformed^18,19^. A greater number of compounds can therefore directly promote ecosystem multifunctionality by increasing the number of ways that microbial communities and abiotic processes transform organic matter^13^. For example, adding compounds to a mixture can increase the variety of unique chemical properties and thus the number of organic compounds that are degraded, subsequently increasing microbial biomass. Second, chemodiversity can indirectly influence ecosystem multifunctionality by changing the physical environment. For example, because compounds vary in their ability to absorb light^20^, the addition of light-absorbing compounds to aquatic ecosystems can reduce photosynthetic rates and limit heat penetration into deeper waters^21,22^. These changes can result in warmer and more nutrient-rich surface waters that accelerate microbial activity and carbon cycling between the land surface and atmosphere^6,23^. For these reasons, the number and characteristics of organic compounds in the environment can provide an integrative measure across multiple ecosystem functions whose importance can be compared directly with other variables traditionally used to explain ecosystem multifunctionality, including biodiversity^9^ and climate^3,8^. However, whether chemodiversity explains a relatively large amount of variation in ecosystem multifunctionality remains untested.

Here we tested the hypothesis that chemodiversity promotes ecosystem multifunctionality by combining a manipulative lab experiment with the largest field survey of these variables to date. We focused on lakes unlike past studies that tested associations between biodiversity and ecosystem multifunctionality in terrestrial ecosystems^8^. Despite covering only 3.7% of the Earth’s surface, the functions of lakes, including carbon cycling and food web production^23,24^, are disproportionately important to many services valued by society, such as climate regulation and food provisioning. We first used a lab experiment to test how changing chemodiversity changes ecosystem multifunctionality, and then tested the importance of this association in natural lakes that span 30**°** of latitude as compared with known drivers of ecosystem multifunctionality, namely biodiversity and physicochemical variables. We defined multifunctionality from a human perspective that values services like climate regulation and food web productivity, and measured chemodiversity in surface waters using ultrahigh-resolution Fourier-transform ion cyclotron resonance mass spectrometry (FT-ICR MS) to identify the diversity of compounds involved in biotic and abiotic reactions. Although the analytical window of FT-ICR MS can omit low (<150 m/z) and high-molecular-mass (>1,000 m/z) compounds that contribute to biotic and abiotic reactions^25^, it is the most sensitive tool available for characterising chemodiversity and captures most of the organic compounds that interact with microbes^26^. We then used the field survey and current theories that synthesise how different types of chemodiversity originate^13,27^ to develop a general understanding of how and where chemodiversity matters for ecosystem multifunctionality. As lakes are more abundant at northern latitudes^28^ that are undergoing large changes in organic matter inputs, we use our results to forecast the consequences of environmental change for chemodiversity and the potential impacts on the benefits that lakes provide to society.

### Chemodiversity is the main driver of ecosystem multifunctionality

To test if ecosystem multifunctionality increases with increasing chemodiversity, we first exposed four levels of biodiversity (1, 2, 3, and 6 bacterial species) under controlled laboratory conditions to three environmental sources of dissolved organic matter (DOM). The three DOM sources naturally varied in chemodiversity (1090, 1745, and 2564 formulae), which closely correlated with indices summarising potential biogeochemical functions, such as defined from the bioavailability, reactivity, and molecular mass of DOM^29^ (Fig. S1). The molecular composition of the three DOM sources was also largely nested (Fig. S2), whereby increasing chemodiversity primarily consisted of the addition of aromatic-like formulae (Table S1). Our fully factorial design allowed us to compare the importance of chemodiversity versus biodiversity for predicting ecosystem multifunctionality from the main effects of chemodiversity and biodiversity and their interaction. Ecosystem multifunctionality was defined using changes in cell abundances, dissolved organic carbon (DOC) concentrations, and total dissolved nitrogen (TDN) concentrations over a 7-day period. We chose these functions because they are important measures of food web productivity and nutrient cycling^30^. Because high microbial activity can increase both DOM and TDN concentration by releasing exudates or transforming particulate organic matter into dissolved compounds, as well as reduce DOM and TDN concentration by assimilating and respiring organic compounds^31^, we considered the absolute change in DOC and TDN concentration as an indicator of microbial activity. We found that these three functions were largely orthogonal to each other in our closed system, thereby representing complementary axes of ecosystem multifunctionality (Fig. S3). We then summed standardized values (*z*-scores) of the change in each of the three functions between day 0 and day 7 into a single index of ecosystem multifunctionality as is common in ecology^1,3^. Larger values of this index reflect higher ecosystem multifunctionality. We tested the main and interactive effects of chemodiversity and biodiversity as factors on multifunctionality using a linear model without any other variables, i.e. only accounting for the experimental design (see Methods).

We found strong experimental support for our hypothesis that chemodiversity enhances ecosystem multifunctionality. Increasing chemodiversity led to higher values of both the multifunctionality index (Fig. 1b) and each individual ecosystem function, with one exception (Fig. S4). Over the range of observed chemodiversity (1090 to 2563 formulae), multifunctionality was estimated to increase from a mean (95% confidence interval [CI]) of -0.90 (-1.49 – -0.32) to a mean of 1.44 (0.10 – 2.77) (Fig. 1a). This mean increase corresponded to an absolute change of 260% (107 – 966%). In support of these findings, DOM was increasingly transformed with increasing chemodiversity. We specifically found ten-times as many compounds were consumed at the highest compared to the lowest level of chemodiversity, whereas the percentage of compounds that were produced in the incubations, such as might arise from cell lysis, was relatively stable across chemodiversity treatments (Fig. S5). Importantly, we found no evidence that chemodiversity was an outcome of greater multifunctionality. The change in chemodiversity, that is, the difference between chemodiversity measured at the end of the incubation in control incubations with no microbes and chemodiversity measured at the end of the incubation in the presence of microbes, was not correlated with multifunctionality (ρ=-0.22, p=0.203). Chemodiversity alone explained 24% of the variation in multifunctionality, which is greater than the effect of biodiversity in many other studies of ecosystem multifunctionality^3,8–10^.

**Figure 1:**
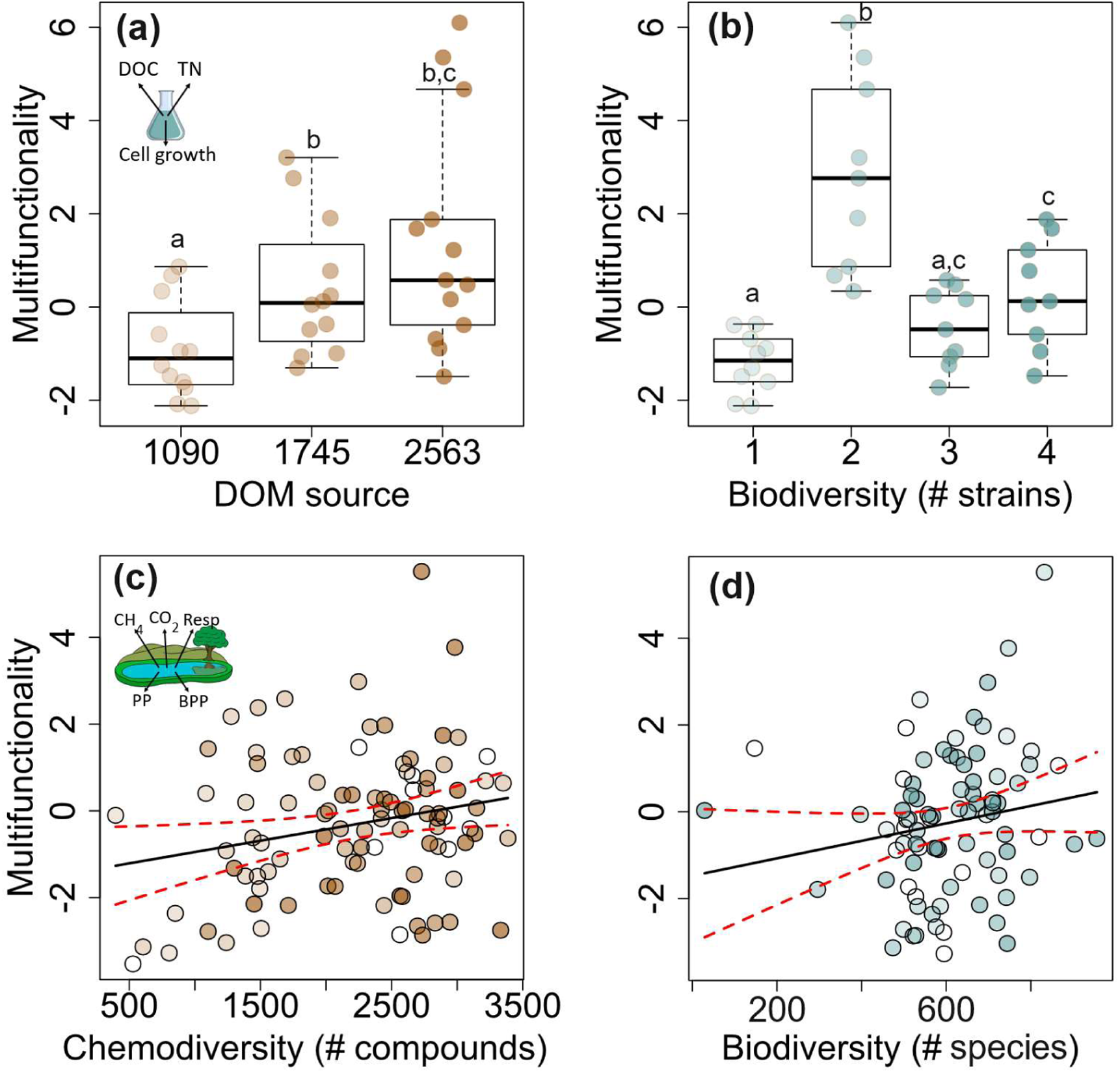
Chemodiversity increases ecosystem function. Multifunctionality increased with **(a)** higher chemodiversity but not with **(b)** higher biodiversity under controlled conditions. Multifunctionality was measured as the sum of *z*-scores of changes in each of cell abundances, and transformation of dissolved organic carbon (DOC) and total dissolved nitrogen (TDN) over 7 days in 36 microcosms represented by individual points. Boxes encompass the inter-quartile range with median denoted by solid horizontal line and whiskers equal to 1.5-times the interquartile range. Different letters indicate statistically significant (p<0.05) differences between factor levels with Tukey’s pairwise comparison tests. Chemodiversity of each experimental source was estimated from the exponential of the Shannon diversity index. In a separate field study, multifunctionality increased with **(c)** higher chemodiversity but not with **(d)** higher biodiversity across 82 European lakes represented by individual points. Multifunctionality was measured in the field study as the sum of *z*-scores of CH_4_ and CO_2_ emissions, total respiration (Resp), maximal net primary production (PP), and bacteria protein production (BPP). Lines in **(c)** and **(d)** are estimated mean associations based on a linear model ± 95% confidence intervals. For all, darker colours indicate higher chemodiversity (brown points) and biodiversity (blue points).

Neither ecosystem multifunctionality (Fig. 1b) nor any individual function increased consistently with biodiversity in our experiment (Fig. S4). Although the two species community had higher multifunctionality, its taxa were included in the six species community (Fig. 1b). Thus, if increasing biodiversity increased ecosystem function, we would expect the six species community to have an equal or higher multifunctionality than the two species community. However, we observed lower multifunctionality with six species than with two species (Fig. 1b). Adding biodiversity can sometimes have no added benefit for ecosystem function, potentially because of competition and redundancy in the functions undertaken by different species^32,33^. In our experiment, we found that the two species community grew faster and caused larger absolute change in DOC and TDN than when its members were grown alongside other taxa in the six species community (Fig. S4), suggesting competitive inhibition in the latter. Most microbial taxa negatively interact when grown together^34^, which reduces their growth relative to expectations from when taxa are grown by themselves^35^.

We also observed an interaction between chemodiversity and biodiversity that further emphasizes the importance of chemodiversity for ecosystem multifunctionality. Higher biodiversity increased ecosystem multifunctionality only at the highest chemodiversity level (Fig. S6). These results suggest that without a sufficient diversity of compounds for microbes to use, increasing biodiversity alone cannot sustain a higher ecosystem multifunctionality. By contrast, in environments with low biodiversity and low chemodiversity, increasing chemodiversity alone can increase ecosystem multifunctionality by enabling the species already present to perform more functions. Because ecosystem multifunctionality-biodiversity relationships^3,8–10,32,36–38^ are mostly studied in terrestrial environments, where chemodiversity is far higher than in aquatic environments^39^, the importance of chemodiversity may have been overlooked.

A potential criticism of our experiment is that because we observed an increase in absolute carbon and nitrogen concentration, changes in ecosystem multifunctionality may have been due to cell lysis rather than microbes transforming chemodiversity. However, both carbon and nitrogen could have come from particulate organic matter as well as nitrogen fixation by some of our study species^40,41^. In support of the former explanation, between 1.6 to 10.6% of the total organic carbon in each treatment was in particulate rather than dissolved form. We also found evidence that nitrogen was incorporated into DOM from other pools than cell lysis. Specifically, we found that a higher proportion of N-containing compounds were produced at higher chemodiversity, and this result could not have been due simply to cell lysis because there was a higher overall consumption of compounds (Fig. S5). Taken together, our results suggest that increasing chemodiversity increases ecosystem multifunctionality by enabling microbes to use and transform substrates in more ways than is achieved by increasing biodiversity alone.

To test the generality of our lab experiment, we sampled surface waters from 101 natural lakes spanning continental Europe (∼30° latitude). As in the lab experiment, the proportion of aromatic compounds increased from 3.1% to 28.4% over the range of chemodiversity observed in the field from 396 to 3387 molecular formulae (linear model, t=6.54, df=100, p<0.001). The chemical characteristics of DOM, including its molecular weight, further suggested that aromatic-like compounds were increasing with chemodiversity like in the lab experiment (Fig. S7). We also measured different functions in the field than in our experiment to test if chemodiversity promoted ecosystem multifunctionality irrespective of the specific functions being measured, that is, if the association between chemodiversity and ecosystem multifunctionality was generalizable. The functions measured in the field were related to carbon cycling (CH_4_ and CO_2_ emissions) and ecosystem metabolism (total respiration, maximum net primary production, and bacterial protein production). We reversed the sign of the measured values for CH_4_ emissions, CO_2_ emissions, and total respiration so that a higher multifunctionality value corresponded to a greater benefit to society^37^ (see Methods). Thus, larger values of these three functions indicated lakes were both emitting and mineralizing less carbon. We then summed *z*-scores of the five functions into a single index as in the lab experiment. We also compared the importance of chemodiversity with the importance of microbial (bacteria, fungi, archaea) and eukaryotic biodiversity estimated from shotgun metagenomic sequencing of environmental DNA in 82 of the study lakes. Our biodiversity sampling detected more taxa than in other large-scale studies^42,43^ with between 97 to 1589 species (median = 610) per lake.

We found a positive association between ecosystem multifunctionality and chemodiversity but not with biodiversity across 82 of the 101 field sites with data for all these three variables. Over the range of chemodiversity (396 to 3387 molecular formulae, median = 2445), multifunctionality was estimated to increase from a mean of -1.27 (95% CI: -2.16 – -0.37) to 0.30 (-0.33 – 0.92) (linear model: t=2.24, p=0.030, df=81; Fig. 1c). This mean increase corresponded to a change of 124% (43 – 144%). Neither biodiversity nor its interaction with chemodiversity influenced multifunctionality (t=1.26, p=0.21 and t=-0.02, p=0.99, respectively, df=81 for both). Much like in our experiment, the effect of biodiversity on multifunctionality was limited by chemodiversity. We found that the effect of biodiversity on multifunctionality only became statistically significant at the highest levels of chemodiversity (Fig. S8). Three of the five measured functions, CH_4_ and CO_2_ emissions and bacterial protein production (BPP), also varied individually with chemodiversity whereas only CO_2_ emissions significantly decreased with biodiversity (Fig. S9). At a molecular-level, these functions were associated with different formulae, potentially reflecting differences in how microbial functions consumed organic substrates. For example, BPP was negatively correlated with generally low H:C formulae (Fig. S10), indicative of consumption^4^, because these formulae were relatively more abundant and thus easier for bacteria to find to build cellular structures than rarer high H:C compounds that may release more energy^4,44^. CH_4_ was positively associated with the proportion of formulae per lake that contained either N or S (ρ=0.24, p=0.006), which are precursors for oxic methanogenesis^45,46^, and CO_2_ negatively correlated with high H:C formulae (Fig. S10), which are preferably respired by bacteria because they provide the highest energy return^4,44^. Chemodiversity alone explained 5% of the variation in multifunctionality, which is lower than in our experiment but we considered more functions than in our experiment and our experiment controlled for environmental noise, e.g. climate. This result is also comparable to, and in some occasions up to five-times higher than, the effect of biodiversity reported in influential field studies explaining variation in ecosystem multifunctionality^8,36^. The choice of functions affects the strength of the relationship between biodiversity and multifunctionality, so may explain some of these differences among studies.

To test further the importance of chemodiversity relative to published estimates of biodiversity, we computed multifunctionality based on the exceedance of data-derived thresholds^37^. This method was specifically designed to facilitate inter-study comparisons^37^. We found that the reduced explanatory power of biodiversity relative to chemodiversity persisted when we estimated ecosystem multifunctionality with this threshold approach (Supplementary Note 1). Our estimated effect of chemodiversity on multifunctionality with this threshold approach was larger than the effect of biodiversity on ecosystem function that was previously estimated from 47 published studies^9^, including freshwaters (Supplementary Note 1). Unlike for biodiversity-ecosystem multifunctionality, where positive associations have been reported in studies measuring plant productivity^3,8^, and rarely in studies measuring macronutrient cycling^33^, the chemodiversity-ecosystem multifunctionality relationship may be less sensitive to the specific functions measured. We found consistent positive associations between chemodiversity and ecosystem multifunctionality across our experimental and field datasets with fundamentally different functions associated with primary productivity and macronutrient cycling. Together, our results from both controlled and natural conditions suggest that the large role of chemodiversity in supporting ecosystem multifunctionality has, to date, been overlooked and must be considered in efforts aimed at delivering benefits from nature to society.

### Recalcitrance underlies positive chemodiversity-multifunctionality associations

Based on theoretical models that predict that the reactivity of DOM varies with its molecular composition and local environmental conditions^13,27^, we hypothesised that the effect of chemodiversity on ecosystem multifunctionality depends on the source of chemodiversity itself. By sampling lakes at similar periods of annual heat accumulation, and with a similar light environment and DOC concentration (see Methods), we found that spatial variation in chemodiversity at a continental scale primarily reflected differences in climate and surrounding land use^16,47^ (Fig. S11). Specifically, chemodiversity strongly increased from southern to northern Europe (ρ=0.40, p<0.001), suggesting that the environment in which chemodiversity occurred and/or its source could explain why chemodiversity varied throughout Europe. Here we extend these two potential explanations (see Supplementary Note 2 for further discussion) to contend that they differentially influence how chemodiversity is used by microbes and thus ecosystem multifunctionality. If chemodiversity was high because of environmental conditions (“environment” hypothesis in Fig. 2a) that slow microbial transformation of DOM, such as lower mean annual temperature (MAT) or lower macronutrient concentrations, we expected that ecosystem functions like microbial respiration and bacterial protein production would also be reduced^48^, resulting in a negative correlation between chemodiversity and multifunctionality. By contrast, if chemodiversity primarily arose because compounds with greater chemical complexity were imported into lakes, we expected these compounds to support a greater number of biochemical transformations and increase ecosystem multifunctionality (“source” hypothesis in Fig. 2a). Complex compounds, such as those with high aromaticity, are abundant in lakes and their degradation by microbes involves many biochemical pathways that increase metabolic activity and biomass formation^4,49,50^, and ultimately multifunctionality. By contrast, simpler compounds are expected to decrease multifunctionality because they are often used to produce energy, which emits CO_2_ as a by-product.

**Figure 2:**
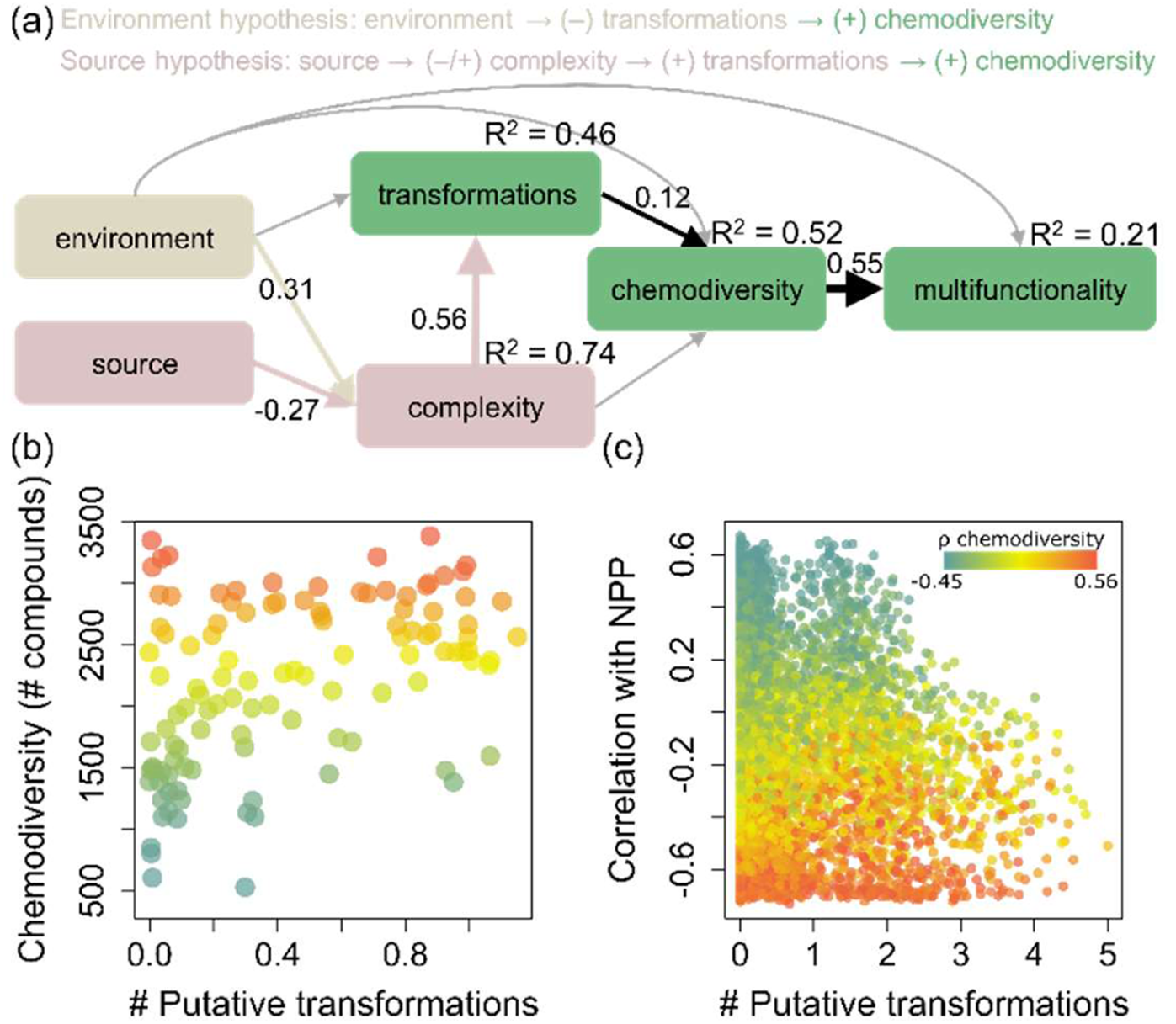
The origin of chemodiversity influences its effect on ecosystem multifunctionality. **(a)** We hypothesized that the positive effect of chemodiversity on ecosystem multifunctionality was caused by environmental conditions reducing the transformation of DOM (“Environment” hypothesis) and sources of DOM producing compounds that could be transformed in a greater number of ways (“Source” hypothesis). + and – symbols show hypothesised cause-and-effect of one variable on another, with either positive or negative links between NPP and complexity possible in the Source hypothesis. We tested these hypotheses using path analysis fitted to 101 lake-level observations. To simplify the path analysis, we created a composite variable “environment” that encompassed 9 measured parameters (see Fig. S13 for effect of each parameter), whereas “source” was equal to NPP in the surrounding catchment of each lake. Complexity and transformations corresponded with the intensity-weighted modified aromaticity index and mean number of putative transformations per molecular formula per lake, respectively. Coloured arrows show statistically significant paths with standardized effects. **(b)** The mean number of transformations per formula per lake was closely correlated with chemodiversity (ρ=0.40, p<0.001). Each dot represents a lake (N=101) coloured for chemodiversity. **(c)** We also correlated the number of transformations that each of 8456 molecular formulae underwent across all lakes with the NPP of the lake catchment. Formula (points) associated with low NPP (i.e., negative value along the *y*-axis) generally underwent more transformations (i.e. larger value along the *x*-axis; ρ=-0.10, p<0.001). Numbers of transformations were weighted by the relative abundance of each formula across lakes. Points are coloured according to their correlation with chemodiversity with low (blue) and high (red).

We tested how both the complexity of DOM and the environment in which DOM was degraded affected microbial transformation of DOM and subsequently the effect of chemodiversity on ecosystem multifunctionality. We estimated DOM complexity in each lake based on an aromaticity index^51^ (see Methods). We expected this index to reflect terrestrial vegetation^16^, which is a main source of lake DOM^40^. Slower growing vegetation tends to invest more in protecting biomass, such as with secondary metabolites^52^, because it faces a relatively greater cost to produce biomass. Compounds involved in plant protection tend to be more complex and have chemical properties, including a higher aromaticity, which require more biochemical reactions to be transformed^14,27^. We expected complexity to influence multifunctionality by determining the extent to microbes can transform DOM and we expected complexity itself to be determined by terrestrial net primary productivity (NPP), i.e., vegetation growth rates, in surrounding catchments. We also considered nine environmental variables (MAT, annual precipitation, water temperature, pH, water colour, lake area, biodiversity, trophic state, and nutrient concentrations) that could influence chemodiversity by changing the diversity of terrestrial and aquatic organic matter inputs, controlling the physical energy required to break chemical bonds, and regulating photo-oxidation^53,54,55^. NPP and the environmental variables were estimated from high-resolution remote sensing data and in-situ measurements (see Methods). We then averaged the number of times molecular formulae could be transformed in each lake using a list of 1255 putative biochemical transformations^19^ (see Methods). If the positive association we observed between chemodiversity and ecosystem multifunctionality arose from the input of complex compounds rather than the limitation of microbial transformations of DOM, we expected NPP would be more strongly associated with chemodiversity than the nine other environmental variables.

Our results indicate that the source of DOM explains how and where a positive chemodiversity-ecosystem multifunctionality relationship can arise. Specifically, we found that the production of complex compounds by low terrestrial NPP (ρ=-0.70, p<0.001, Fig. S12a) correlated with both greater chemodiversity (ρ=0.56, p<0.001; Fig. S12b) and a greater number of average transformations that formulae could undergo in each lake (ρ=0.45, p<0.001, Fig. S12c). Consequently, chemodiversity correlated positively with the number of transformations per formulae (ρ=0.40, p<0.001, Fig. 2b). Correlations between chemodiversity and each of complexity, NPP, and the number of biochemical transformations are predicted by the source hypothesis. By contrast, all the environmental variables had weaker associations with chemodiversity, though some were statistically significant (Table S2). Together, our results indicate that the production of complex compounds by low NPP increases both chemodiversity and the number of biochemical transformations that each compound can support.

We synthesised the observed correlations among chemical complexity, biochemical transformations, and chemodiversity to propose a new general understanding of ecosystem multifunctionality. This new understanding was that the positive effect of chemodiversity on multifunctionality arises because of the production of complex compounds by plants and environmental conditions that favour those compounds within DOM pools. We found further support for this framework with a path analysis explicitly testing presumed causal pathways. The path analysis modified our original hypotheses to incorporate additional paths identified by tests of model structure (Fig. S13). With this analysis, we found that effects related to the production of organic compounds (i.e. NPP) influenced chemodiversity by changing DOM complexity and subsequently the number of biochemical transformations it can receive (Fig. 2a). Although environmental variables (i.e., water colour and MAT) also influenced DOM complexity, the direction of these effects suggested that they were associated with reduced in-lake primary productivity and an increase in the relative importance of terrestrial DOM (Fig. S13). This interpretation was further supported by the direct negative effect of chlorophyll *a* concentrations on chemodiversity (Fig. S13).

To test further if ecosystem multifunctionality was enhanced by chemodiversity because of the properties of complex compounds associated with low terrestrial NPP, we analysed the composition and putative source of all 8456 molecular formulae identified in the field survey. We first identified formulae whose relative abundances across our study lakes were correlated with terrestrial NPP of the surrounding catchment. Positive and negative correlations suggest formulae likely originated from high and low terrestrial NPP, respectively. We compared these correlations with the average number of biochemical transformations that each formula was predicted to undergo in each lake^19^. This analysis allowed us to infer the origins of compounds that can be highly transformed by biotic and abiotic reactions, which contribute to higher ecosystem multifunctionality. Compounds that were associated with low NPP appeared more transformed (t=-8.19, df=6639, p<0.001), and positively correlated with chemodiversity (t=8.50, df=6639, p<0.001; warmer colours in Fig. 2c). These compounds were more complex, on average, with a mean (± standard error) aromaticity index of 0.52 (±0.17) compared to 0.32 (±0.24) in the entire dataset (Wilcoxon rank sum test, W=5.4 × 10^6^, p<0.001, Fig. S14). These results further supported the hypothesis that complex compounds can be degraded in more ways and support higher multifunctionality. We then selected molecular formulae that most increased in abundance with decreasing NPP, that is, were in the top 10% of negative associations based on Spearman rank correlations. We found that these formulae could support, on average, 1.4 times more known putative biochemical transformations than formulae that were weakly associated with NPP, i.e. lowest 10% of negative associations (t=5.80, df=860, p<0.001), and 1.8 times more transformations than formulae strongly positively associated with NPP, i.e. top 10% of positive correlations (t=8.00, df=1090, p<0.001). Together, these results demonstrate that the effect of chemodiversity on multifunctionality emerges from the properties of individual formulae.

We expect chemodiversity will be less important for ecosystem multifunctionality in environments where microbes cannot access and transform large quantities of organic matter. For example, DOM concentrations are typically >5 mg C L^-1^ in rivers, most lakes, wetlands, or mangroves^19,47,48,56^, but <1 mg C L^-1^ in the deep ocean^29^ or in large (surface area >500 km^2^) deep lakes with centennial residence times^57^. At these concentrations, individual compounds are more likely to be too dilute to interact with microbes^52^, irrespective of the diversity of compounds in the environment, and may result in a shallower slope between chemodiversity and ecosystem function. In support of this expectation, we found that compounds transformed in lakes reacted less frequently when found in the deep ocean. We identified 2745 formulae shared between our study lakes and a representative deep ocean sample^18^, which is likely an overestimate because we cannot differentiate structural isomers. We then calculated the number of times that each formula was transformed in each sample using the established list of putative biochemical transformations^19^. We found that these formulae underwent, on average, 0.08 (range: 0 to 1.32, standard deviation = 0.17) putative transformations in lakes but no transformations were detected in the deep ocean. Given that many of the detected transformations are ubiquitous, such as methylation, a likely explanation for why we did not find them in the ocean sample is because their substrates occur at concentrations lower than our femtomole detection limit^58^. Such concentrations would be several order of magnitude beneath concentrations of the most limiting organic compounds^59^. This observation further suggests that chemodiversity will have a greater influence on ecosystem multifunctionality where individual compounds occur at high enough concentrations to support many biochemical transformations.

### Chemodiversity and ecosystem multifunctionality under global change scenarios

NPP is expected to increase because of rising atmospheric CO_2_ concentrations^60^, and our results suggest that this increase could reduce the chemodiversity of lakes in ways that are relevant to global carbon budgets and climate modelling efforts. To predict how the resulting changes in chemodiversity will influence ecosystem multifunctionality, we first modelled the value of each ecosystem function that varied with chemodiversity in our field survey (Fig. S9). By adding chemodiversity to statistical models of CH_4_ and CO_2_ emissions and BPP that contained predictors identified as important in previous large-scale studies^23,24^, we increased explained variation of each function by up to 50% as compared with past studies (Table S4). We achieved the large increase in explained variation despite being unable to measure some prominent drivers of ecosystem metabolism, such as phosphorus concentrations^61^. Importantly, based on the properties of linear models, additional predictors are unlikely to change the effect of chemodiversity on individual functions and will only further reduce unexplained variation. We then projected our models for each ecosystem function to all European lakes >10 ha (see Methods). For chemodiversity, we generated a European-scale raster with the model used in the path analysis (Fig. S13). This analysis further validated our study because we found that the extrapolated estimates were within bounds of global-scale studies^23,24^; no European-only studies exist for comparison. Overall, we estimated that European lakes emitted, on average, 22.7 Tg C yr^-1^ as CH_4_ (95% CI 11.2 – 32.0; model R^2^ = 0.46) and 27.8 Tg C yr^-1^ as CO_2_ (95% CI: 9.9 – 37.4; R^2^ = 0.22) in 2019. Despite these emissions potentially being overestimated because they only considered summertime fluxes^62^, they corresponded to 7% and 19% of estimated global lake CH_4_ and CO_2_ emissions^23,24^, respectively, and Europe contains ∼20% of the world’s total number of lakes^63^. We also estimated that European lakes fixed an average of 1.8 Tg C yr ^-1^ (95% CI: 1.0 – 3.3; R^2^ = 0.45) in 2019 into microbial biomass, that is, from BPP.

To forecast the consequences of changes in NPP for European lakes, we input NPP estimated in the year 2100 by Earth system models^64^ into the linear model that predicted chemodiversity (Table S3) and then used the predictions to forecast each ecosystem function (Table S4). We focused solely on changes in NPP, holding other variables constant, because NPP was the best individual predictor of chemodiversity (Table S2) and had a clear mechanistic link to chemodiversity (Fig. S13). We used two shared socioeconomic pathways (SSPs) representing likely (SSP2) and worst case (SSP5) scenarios for future changes in atmospheric CO_2_ concentrations. We found that future increases in NPP alone will reduce chemodiversity in 99.1% of lakes, with the median chemodiversity decreasing by 13% and 24% under the SSP2 and SSP5 scenarios, respectively (Fig. 3a). When summed across lakes, we estimated that the change in chemodiversity will cause lakes to emit an additional mean 28.4 (95% CI: 3.3 – 166.1) and 32.2 (95% CI: 14.0 – 231.2) Tg C yr^-1^ as CH_4_ under SSP2 and SSP5 scenarios, respectively (Figs 3b, S15). Similarly, the changes in chemodiversity will cause lakes to emit an additional 3.7 (95% CI: 1.3 – 5.1) and 4.0 Tg C yr^-1^ as CO_2_ (95% CI: 1.3 – 5.2 Tg C yr^-1^) under SSP2 and SSP5 scenarios, respectively (Figs 3c, S15). Microbes will, however, fix 4.3 (95% CI: 1.7 – 10.2) and 4.4 Tg C yr^-1^ (95% CI: 1.7 – 12.6 Tg C yr^-1^) more carbon into biomass under SSP2 and SSP5 scenarios, respectively (Figs 3d, S15). Together, our estimates suggest that European lakes may emit, on average, 1.6-times more carbon by 2100, mostly as CH_4_, because of NPP-associated declines in chemodiversity. Although our estimates are higher than previous estimates of future CH_4_ emissions by lakes in absolute terms by 50%, they are comparable in terms of relative change, i.e. 73-84% in our study versus 58-88%^62^. The difference in absolute terms is likely due to the lack of winter measurements in our dataset as lower microbial activity in colder weather typically reduces greenhouse gas emissions^62^. The similarity in relative terms instead emphasises that chemodiversity is a necessary predictor of CH_4_ emissions.

**Figure 3:**
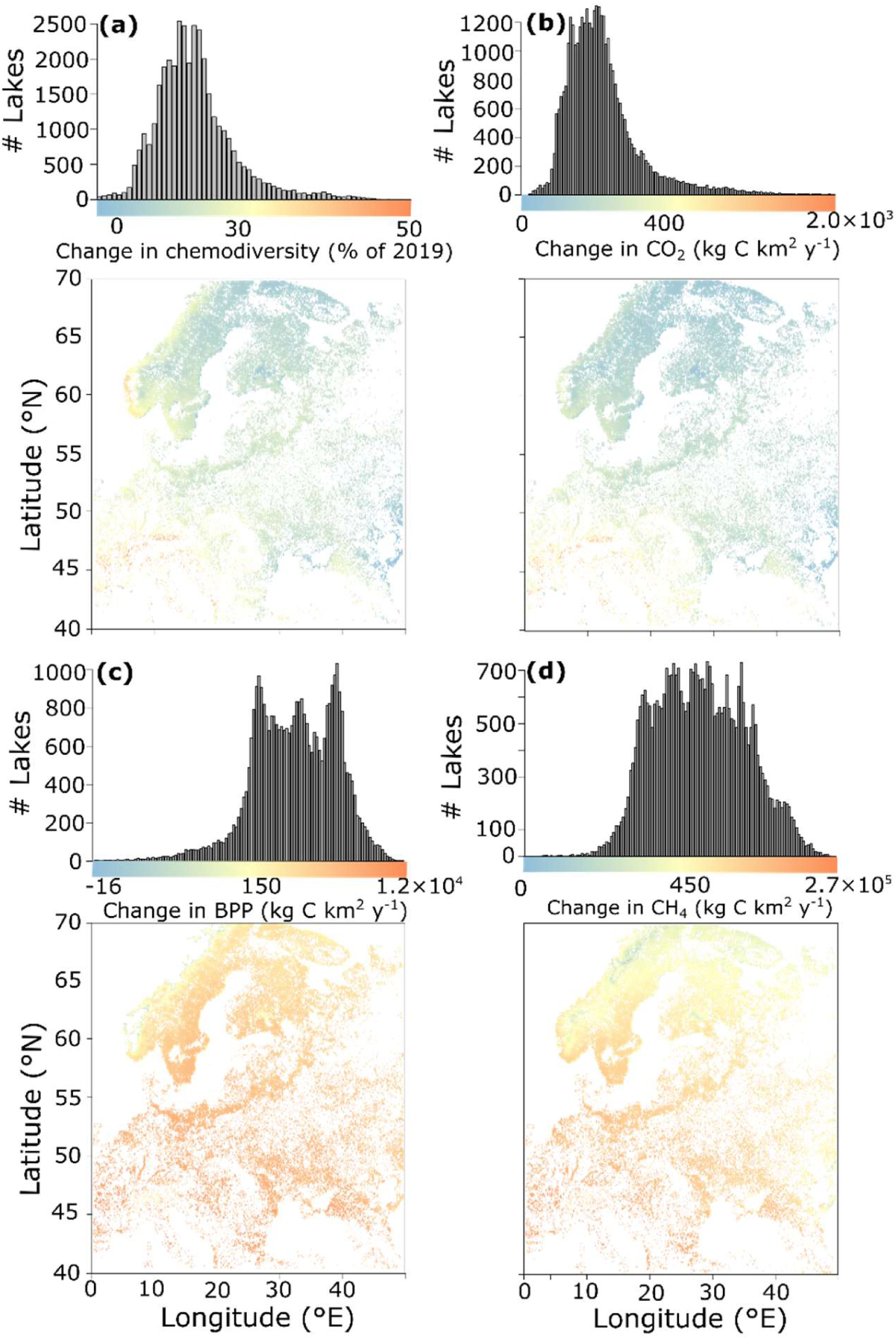
Lakes will emit more carbon by 2100 due to continental-wide declines in chemodiversity. We estimated **(a)** chemodiversity in 2100 while holding all other variables at 2019 levels for all European lakes >10 ha in surface area (R^2^ for chemodiversity model = 0.36). Values are the change in chemodiversity compared to 2019. Using future chemodiversity, we then estimated mean changes in **(b)** CO_2_ emissions **(c)** bacterial biomass production, and **(d)** CH_4_ emissions from pelagic surface waters between 2019 and 2100 using linear models (R^2^ = 0.22, 0.45, and 0.46, respectively; Table S4). For all, points indicate N=175,500 individual lakes. Histograms indicate the number of lakes found with corresponding changes. 95% confidence intervals for each function are given in Fig. S15.

We forecasted that lakes in southern Europe, and in the south of the Scandinavian peninsula, will experience the largest changes in ecosystem function (Fig. 3). These lakes are located near large population centres and, therefore, provide opportunities to improve management of their surrounding catchments in ways that can enhance the benefits that European lakes provide to society. For example, our results suggest that chemodiversity and subsequently ecosystem multifunctionality can be enhanced by promoting slow growing vegetation, such as many Fabaceae^65^. Slow growing vegetation produce large numbers of complex polymeric compounds, which undergo many biochemical transformations when leached into water from litterfall and soils^17^. They also have higher wood density^66^, that can increase long-term carbon storage on land^67^, resulting in a win-win for land and water.

### Conclusions

Here, we provided experimental and large-scale observational evidence that increasing chemodiversity delivers beneficial ecosystem functions, especially a stronger carbon sink capacity corresponding with greater bacterial biomass and lower greenhouse gas emissions. We then demonstrated that the ways in which ecosystems deliver these functions depends on the molecular properties of DOM, namely the capacity of individual compounds to undergo biochemical transformation. The molecular properties of compounds ultimately depend on sources of DOM and external environmental conditions, such as their concentrations, as found in many inland waters, wetlands, and mangroves^19,47,48,56^. Here we add to this understanding by showing the importance of these properties for ecosystem multifunctionality and their potential to explain ecosystem-specific differences in chemodiversity–multifunctionality relationships. Our findings also suggest that chemodiversity and its effect on multifunctionality can be altered across different aquatic ecosystems by manipulating sources of DOM within surrounding catchments. Finally, our study suggests that accounting for the strong association between NPP and chemodiversity can improve integration between terrestrial and aquatic ecosystem models^5^. For example, because chemodiversity is strongly predicted by NPP, which itself is predictable at high temporal and spatial scale from bioclimatic data, we can readily incorporate the effect of chemodiversity on greenhouse gas emissions into existing carbon budgets. More generally, our study demonstrates that sustaining the benefits that lakes provide to people will require management that extends beyond shorelines and considers the chemodiversity exported from land.

## Acknowledgments

We are grateful for help with field work from Samuel Cottingham, Carolyn Ewins, Sophie Guillaume, Eleanor Sheridan, and Samuel Woodman. We thank Yi Zhang and Caroline Kemp for laboratory assistance, Del Hawtin, and Simon Hoer for logistical support, and Katrin Klaproth for assistance with FT-ICR MS measurements. We thank Jim McGeer for providing the DOM sources and Thomas Scheuerl for providing the bacterial strains used in the experiment. This work was funded by H2020 ERC Grant sEEIngDOM #804673 to AJT and the University of Cambridge Harding Distinguished Postgraduate Scholars Programme.

## Author contributions

AJT and JAF designed the study. JAF performed the field and associated lab work with input from AJT and TD. SRS and JAF performed the laboratory experiment. JAF analysed the data with input from AJT. JAF and AJT wrote the manuscript with input from TD and SRS.

## Data availability

All data are available within the manuscript or upon reasonable request.

## Materials and methods

### Laboratory experiment

To test if multifunctionality varies with chemodiversity, we incubated bacterial communities of four biodiversity levels with three different levels of chemodiversity. The four biodiversity levels were 1) *Pseudomonas fluorescens* SBW25 2) *Chryseobacterium* sp. and *Sphingobacterium* sp. isolated from natural communities that degrade DOM^68^; 3) *Rahnella* sp., *Raoultella* sp., and *Pantoea* sp. from the same natural communities^68^; 4) all six microbes together. Before starting the experiment, we grew each microbial isolate individually from a frozen archive for five days in R2A media. A control of filtered (0.2 µm, Sterivex-GP, MilliporeSigma, USA) sterile water was included alongside each treatment and indicated no contamination as measured with flow cytometry (see below). For chemodiversity, the three treatment levels were obtained from reverse-osmosis-concentrated water^69^ of a temperate wetland (Luther Marsh, Ontario, Canada; 43°57’43”N, 80°23’50”W), boreal lake (Clearwater Lake, Ontario, Canada; 46°22’11”N, 81°03’04”W), and Arctic river (Cameron River, Northwest Territories, Canada; 62°29’33”N, 113°32’ 48”W). Using MilliQ water, we diluted each reverse-osmosis concentrate to a final dissolved organic carbon (DOC) concentration of 10 mg C L^-1^, as measured with a TOC-TNM-L analyser (Shimadzu, Japan), and filled fifteen 250 mL pre-combusted (4 hr at 550°C) Erlenmeyer flasks with 100 mL of each water source (n = 3 replicates for each of 4 biodiversity levels with 3 controls). Before inoculating the DOM sources with each microbial community, we estimated strain-specific cell number with a CLARIOstar fluorometer (BMG Labtech, UK) after 30 min of staining with SYBR Green (10,000× stock solution, Invitrogen, USA) diluted to a 10× final concentration. Based on these data, we diluted an unstained aliquot of each strain to an expected 10,000 fluorescence units and mixed them to form the four communities. This normalisation created a substitutive design that ensured each community had a similar abundance of DNA and thus potential activity. Then, we added 1 mL of each of the four microbial communities to each chemodiversity level in triplicate and incubated the flasks at ca. 20°C for seven days in the dark without shaking.

### Measuring chemodiversity

We estimated chemodiversity in the experiment from solid-phase extraction of DOM following a standard protocol^70^. Briefly, 80 mL of each incubation flask was acidified to a pH of 2 by adding HCl, passed through 1 g of a polar styrene-divinylbenzene polymer (Bond Elut PPL, Agilent, USA), and eluted with 3.5 mL of ultrapure methanol (LC-MS LiChrosolv, Merck, Germany). We diluted the extracts to a final concentration of 2.5 ppm DOC in a 1:1 (v:v) methanol:water solution before injecting 100 µL into a 15 Tesla Solarix FT-ICR MS (Bruker Daltonic, Germany) with direct infusion electrospray ionization in negative mode. We collected 200 scans in duplicate for each sample and calibrated the spectra using the DataAnalysis software (Bruker Daltonic, Germany). Masses ranging from 150 to 1000 m/z were exported, and we assigned molecular formulas using the online platform ICBM-OCEAN^71^. We used the N, S, P rule, deleted all singletons, and used isotopic confirmation to accept/reject a formula. We limited formula attributions to C_1-100_H_2:200_O_0-70_N_0-3_P_0-2_S_0-2_ with a tolerance of 0.2 ppm and retrieved a total of 3873 individual molecular formulas. We computed chemodiversity as the exponential of the Shannon diversity index using all assigned molecular formulas^72^ and the summed normalised peak intensity per sample. This index represents the diversity in a community when all formulae are equally common, so corrects for variation in evenness that can bias raw counts^73^ of formulae number. We also computed the ratio of hydrogen to carbon atoms in each compound as an indicator of bioavailability, as described elsewhere^44^. The three DOM sources were largely nested with one another with 93% of formulae present in the Arctic river (low chemodiversity) also present in the boreal lake (medium chemodiversity), 70% of boreal lake formulae present in the temperate wetland (high chemodiversity), and 80% of formulae present in the Arctic river present in the temperate wetland (Fig. S2). The large number of shared formulae among treatments confirmed that sources with higher chemodiversity contained the vast majority of formulae present in sources with lower chemodiversity, i.e. were nested. Additionally, the molecular composition of the DOM used in the experiment differed as much from the lakes sampled in the field study as the lakes differed among themselves (Fig. S16).

### Ecosystem multifunctionality in the laboratory experiment

We estimated the multifunctionality of each replicate community in the experiment by measuring carbon consumption, nitrogen consumption, and bacterial growth rate. At the start and end of the incubation, we measured the DOC and TDN concentrations of each flask using a Shimadzu TOC-TNM-L analyser. Carbon and nitrogen consumption were estimated as the change in concentration between the start and end of the incubation. To measure growth rate, we determined the difference in cell abundance between the start and end of the incubation using flow cytometry. Briefly, we removed 800 µL of water from each flask at the start and end of the experiment. Samples were incubated with 200 µL of 25% glutaraldehyde in the dark at 4°C for 30 min before being preserved at -80°C. Within one week, we thawed the samples, added SYBR Green at a 10× final concentration (10,000× stock solution, Invitrogen, USA), and incubated the samples overnight at 4°C. Then, we injected 100 µL of each sample into a flow cytometer (Attune NxT, Thermo Fisher Scientific, USA) at a rate of 250 µL per minute and used the same instrument settings for all samples. We optimised the gating for each DOM source based on stained/unstained samples using FlowJo v10.9 (BD Biosciences, USA) and used the same gating for each community. Each function was standardized to a mean of zero and standard deviation of one to obtain a *z*-score, which we summed across the three functions to estimate multifunctionality^8^.

### Selection of field study sites

We selected lakes to sample across Europe ensuring there were no systematic biases in DOC concentrations and water chemistry across our latitudinal gradient. Using the EU’s Water Information System for Europe (WISE) database version 4^74^, we subsampled all entries labelled as “lakes” for which DOC was reported in mg C L^-1^. We also selected entries with a total organic carbon (TOC) concentration equal or lower than 5 mg C L^-1^ to include more lakes with low DOC values, which were underrepresented in the WISE database despite being known to be the most common on Earth^75^. From the 2169 resulting entries, we removed 256 entries with a DOC/TOC concentration >30 mg C L^-1^, which is outside the expected values in lakes^75^ and likely represent measurement error or atypical lakes, and 1595 duplicated measurements. We grouped the remaining 318 lakes into five bins in 5 mg C L^-1^ intervals: <5 mg C L^-1^, 5-10 mg C L^-1^, 10-15 mg C L^-1^, 15-20 mg C L^-1^, >20 mg C L^-1^ and six 5° latitudinal intervals: <45°N, 45-50°N, 50-55°N, 55-60°N, 60-65°N, >65°N. The last two DOC/TOC bins (i.e. >15 mg C L^-1^) were absent from the first two latitudinal bins. Therefore, we removed all 47 entries with DOC/TOC concentrations >15 mg C L^-1^. In a global analysis^75^ of lake DOC concentrations, 93% of lakes were predicted to have DOC concentrations <15 C L^-1^, so this filtering minimises potential sampling bias while ensuring we still represent the vast majority of lake DOC concentrations. To standardise the sites further, we considered total nitrogen (TN) and total phosphorus (TP) concentrations of all entries. Six entries lacked TP values, so we discarded all but one, which was located between 50-55°N where entries were sparse. Only 10 entries had TP concentrations higher than 0.2 mg L^-1^, and 21 entries had a TN value higher than 1 mg L^-1^, so we removed these to avoid large differences in lake trophic levels.

After reducing the number of entries to lakes with similar water chemistry, entries were selected based on their distance from one another. Therefore, we discarded 28 entries from eastern Poland, one in Latvia, and two in Hungary as they were logistically challenging to reach. We similarly removed 175 entries situated in the United Kingdom and western France. We then inspected each remaining entry individually using Google Earth satellite imagery, removing 86 sites that were dammed or visibly artificial. This process removed all entries in the WISE database situated south of 40° N and north of 69°N. To extend the latitudinal gradient, we manually added eight sites in southern Italy (<40°N) and seven in northern Norway (>69°N) without knowledge about their DOC/TOC or trophic state. None of the manually chosen lakes were dammed or visibly artificial. This selection process resulted in a total of 109 lakes. During sampling, severe weather conditions, unexpected fences around lakes, or equipment failure prevented us from sampling eight lakes, and one site in Poland was substituted for a lake for which no DOC data were available. Ultimately, we sampled 101 lakes spread over 31 latitudinal degrees. There was no statistically significant correlation between latitude and DOC concentration at the time of sampling (Spearman correlation, ρ=0.05, *p*=0.57, Fig. S17).

### Timing of field sampling

The order in which lakes were sampled was chosen to minimise differences in annual heat accumulation and water temperatures across the latitudinal gradient. Sampling dates were originally chosen to minimise differences in growing degree days (GDD) among lakes. GDD represents the difference between a given temperature threshold at which primary production commences and mean daily temperature^76^. Here we used a threshold of 10°C as is classically employed^76^. To compute GDD, we used the rnoaa package^77^ in R v.4.1.1^78^ to retrieve all weather stations within 50 km of the 101 sampled lakes from the US National Oceanic and Atmospheric Administration National Climatic Data Center database. We determined the median GDD for each day of the year based on temperature data collected from the last 10 years. For years with 366 days, the GDD on the 29^th^ of February was added to the GDD from the 28th of February. Cumulative GDD (cGDD) – representing the sum of GDD from the 1 January until a given date – were obtained by sequential addition for each day of the year for each lake. The computed cGDD indicated that the growing season commenced (i.e. cGDD >10) as early as the 1 January at the southernmost site and ended (i.e. cGDD reached its highest values and no longer increased) as early as the 10 October at the northernmost site.

We then selected sampling dates to survey lakes during similar periods of annual heat accumulation. To find the optimal sampling day, we calculated the variation in cGDD along the latitudinal gradient as the ratio between the 3rd and the 1st quartiles of lake-level cGDD, assuming we started sampling the southernmost site and moved northwards to the closest site at a pace of one lake per day. By starting sampling on the 8 April 2019, we could minimise the variation in cGDD across the latitudinal gradient. Overall, the observed maximal difference between the 5° latitudinal bins was 194.4 GDD, which was much less than the 368 that was originally expected based on the historical data (Fig. S17). This value corresponded with sampling the warmest site 10 days later than the coldest site, during which temperatures remained above 20°C. We further measured underwater irradiance at 1 m depth in 60 lakes across the latitudinal gradient using a HOBO light pendant (UA-002-64, Onset, USA) to test for additional biases in irradiance. Pictures representing the latitudinal gradient are presented in Fig. S18.

### Dissolved organic matter sampling

At each lake, we collected 10 L of surface water in an acid-washed plastic carboy. We sampled from the point with maximum water column depth or the point equidistant to all shorelines when no bathymetry information was available. We expected these locations would be the best mixed in the epilimnion and most representative of the lake water quality. Directly after sampling, 1 L of lake water was passed through a pre-combusted (450°C, 4 hours) glass fibre filter (0.5 µm, Macherey-Nagel, Germany), and the filtrate dispatched into three 40 mL pre-combusted amber borosilicate vials and one 500 mL pre-combusted amber borosilicate bottle. Once on land, we acidified the samples to a pH of 2 by adding HCl. We kept the filtrates in the dark, and either on ice or at ca. 4°C for up to 15 days before shipping them to Cambridge, UK. In the laboratory, we stored the samples at 4°C for up to a week and determined the DOC in each 40 mL vial using a Shimadzu TOC-TNM-L analyser (Shimadzu, Japan). In line with regulatory standards^79^, we averaged DOC concentrations from the three replicates when the standard deviation was within 10% of the mean and averaged the two closest values otherwise. We estimated chemodiversity as described for the laboratory experiment but solid-phase extracted 500 mL of water instead of 80 mL and retrieved a total of 8456 formulae. We also evaporated 10 µL of extract and redissolved it in 10 mL of Milli-Q water acidified to a pH of 2 using HCl to determine the DOC concentration in the extract and corresponding extraction efficiency (mean = 62%, SD =15%).

To determine if chemodiversity promoted multifunctionality because it could undergo more biochemical transformations, we also estimated the number of times that each molecular formula could be biochemically transformed using an established approach^19^. Fudyma *et al.*^19^ compiled 1255 pairs of molecular formulae associated with commonly observed biotic and abiotic transformations, such as enzymatic reactions that add or subtract CH_2_ groups and physical condensation or dehydration, respectively. We considered such a transformation to have occurred if we detected each pair of formulae from a reaction in a sample.

We estimated the complexity of each compound using an aromaticity index (AI_mod_)^51^. A molecule is considered more complex if it possesses more three-dimensional structures and aromatic rings, which are both characterised by the presence of double bonds. However, atoms such as nitrogen can be linked to carbon with double bonds without necessarily forming aromatic rings. By using AI_mod_, we considered both the number of double bound equivalents (DBE), which approximates the saturation of each molecular formula, and the number of heteroatoms, which provides insight into how carbon is bound. The number of DBE and AI_mod_ were computed within the ICBM-OCEAN pipeline using equations described by Koch & Dittmar^51,71^.

### Biodiversity sampling

We estimated biodiversity by sampling environmental DNA in 82 of the study lakes. During sampling, we passed 1L of water through a 0.2 µm filter cartridge (Sterivex, Millipore, USA). The cartridges were stored at -20°C until DNA was using an established protocol^80^. Briefly, filter cartridges were opened in a sterile environment, and we placed the filters in a cryotube containing silica and zirconia beads (3.0, 0.7, and 0.1 mm diameter). We then vortexed the tubes at 2,850 rpm for 15 min to detach cells from the filter. To ensure DNA was released into solution from lysed cells, we added 0.6 mL of phenol-chloroform-isopropanol (25:24:1), 0.6 mL of 5% cetrimonium bromide, 60 µL of 10% sodium dodecyl sulfate, and 60 µL 10% N-lauroylsarcosine before vortexing the tubes a second time at 2,850 rpm for 15 min. We then centrifuged the tubes at 16,000 g for 15 min at 4°C and collected the supernatant. To remove phenol residues, we added 0.6 mL of chloroform-isopropanol (24:1) to the supernatant, gently mixed the samples by inversion, and centrifuged at 16,000 g for 10 min at 4°C. We again collected the supernatant (ca. 0.6 mL) and added 1.2 mL of polyethylene glycol with 1.6M sodium chlorine to precipitate the DNA overnight at 4°C. The following morning, we centrifuged the samples at 17,000 g for 90 min at 4°C to obtain a DNA pellet that we washed with ice-cold (-20°C) 70% ethanol. The pellet was dried at 37°C for 10 min before being dissolved in 50 µL of ultrapure water (Qiagen, Germany). We used the same protocol to extract DNA from a ZymoBIOMICS™ Microbial Community Standard II (Log Distribution) (Zymo Research, USA) and ultrapure water (Qiagen, Germany) that served as positive and negative controls, respectively. The positive control indicated no external contamination, and no DNA was retrieved from the negative control.

We prepared shotgun metagenomics sequencing libraries with the NEBNext Ultra II FS kit (New England Biolabs, USA) following the manufacturer’s instructions. Briefly, DNA was digested for 15 minutes to obtain end polished and A-tailed fragments of ∼350bp. Then, we attached Illumina adaptors by incubating the fragments at 37°C for 15 min in presence of a ligation enhancer. We cleaned the samples using AMPure XP beads (Beckman Coulter, USA) and attached Illumina-specific indexes (New England Biosciences, USA). We purified the product using the AMPure XP beads^81^, quantified DNA concentration on a Qubit fluorometer (ThermoFisher, USA), and pooled libraries at equimolar concentrations before sequencing on a NovaSeq 6000 PE 250 bp (Illumina, USA) at Novogene (Cambridge, UK), who performed demultiplexing and checked read quality (Q30 ≥ 75% and Q20 = 100%). We obtained a similar coverage for all samples (1^st^ and 3^rd^ quartiles: 4.35×10^7^ and 6.70×10^7^ reads). We assigned taxonomy to the reads using Kraken2^82^ and a database containing bacteria, eukaryotes (including fungi), and archaea. We used Bayesian Reestimation of Abundance with Kraken to obtain read counts at the genus level^83^ and estimated biodiversity as the exponential of the Shannon index as for chemodiversity.

### Ecosystem functioning in the field survey

At each field site, we measured three functions related to ecosystem metabolism (maximum primary production, total respiration, and bacterial protein production) and two functions related to carbon cycling (CH_4_ and CO_2_ emissions) from the lake surface.

Primary production and respiration were obtained based on differences in dissolved O_2_ as previously described^84^. First, we filled six glass serum bottles with 20 mL of lake water. All bottles were crimped gastight without air bubbles. Two bottles were processed directly, two bottles were incubated in the dark for 24 hours, and the last two bottles faced a cold light (5000K) lamp and received ca. 34,000 lux for 3 hours. For each bottle, we measured the dissolved O_2_ concentration using optical fibre sensors (Presens, Germany). We recorded dissolved O_2_ concentration every second for approximately five minutes until we reached a plateau. Then, we averaged the last 30 values of the plateau to infer the dissolved O_2_ concentration inside the serum bottles. Primary production was estimated as the difference in dissolved O_2_ between the start of the incubation and after exposure to light. Respiration was the difference in dissolved O_2_ between the start and after 24 hours of dark incubation. We averaged duplicates to obtain one value per lake.

Bacteria protein production (BPP) was estimated based on ^3^H-leucine incorporation^85^. Directly after sampling, we filled six tubes (Eppendorf, Germany) containing 17nM of ^3^H-leucine with 1.5 mL of lake water. We previously added 300 µL of trichloroacetic acid to three tubes that served as blanks. All tubes were immersed inside the lake in a dark container at ca. 0.5 m depth for exactly 1 hour. We terminated the incubation by adding 300 µL of trichloroacetic acid and preserved the samples on ice until we reached the lake shore – the journey time varied from 1 to 30 mins. Then, we centrifuged the samples for 10 min at 16,000 g, removed the supernatant, washed the pellets with 1 mL of 5% trichloroacetic acid, and centrifuged again for 10 min at 16,000 g. We removed the supernatant and let the pellets air-dry for up to 10 min. We added 1 mL of scintillation cocktail (HiSafe, Perkin Elmer, USA) and measured counts per minutes with a portable liquid scintillation counter (Triathler, Hidex, Finland). To obtain the final BPP, we converted counts per minutes into degradation per minutes, subtracted the value of the mean of the corresponding blanks and expressed the results in mg C L^-1^ day^-1^ as previously described^86^. We averaged the values from the three replicates when the standard deviation was within 10% of the mean and averaged the two closest values otherwise.

CH_4_ and CO_2_ emissions were measured using the floating chamber technique^87^. We used a chamber with a 0.07 m^2^ perimeter and 0.002 m^3^ volume that was covered in aluminium tape to prevent heating. We connected the chamber to an ultraportable greenhouse gas analyser (Los Gatos Research, USA) that uses off-axis integrated cavity output spectroscopy using gastight PVC tubing. Before starting any measurements, we let the chamber equilibrate with the atmosphere by waiting until the CH_4_ and CO_2_ curves plateaued. Then, we placed the chamber on the water with 3 cm of the chamber immersed underwater. We monitored the concentration of CH_4_ and CO_2_ inside the chamber every second for 15 mins. We estimated fluxes as the linear slope between the gas concentration and time^87^. We repeated the process (atmospheric equilibration and measurement) twice, and a third time if the two fluxes differed by more than 10%. We then averaged the slopes to estimate CH_4_ and CO_2_ fluxes.

To calculate a multifunctionality index that reflects human values of nature, we assessed the societal benefit of each measured function^1^. We considered that society benefits from higher primary production because it can fuel food web productivity and fix atmospheric carbon into biomass. Similarly, larger BPP benefits society because it increases the amount of carbon in the food web and can store carbon for longer inside lakes. However, higher greenhouse gas (CO_2_ and CH_4_) emissions negatively impact society due to their effect on climate. Higher microbial respiration similarly consumes the carbon stored in lakes and emits CO_2_ into the environment, some of which will escape to the atmosphere. We therefore multiplied each of CO_2_ emissions, CH_4_ emissions and respiration rate by -1 to ensure that they had the same directionality of their societal benefits as primary production and BPP, that is, all functions benefited society more as they increased in value. This decision seemingly contrasted with the lab experiment where we calculated absolute values because both increases and decreases in DOC and TDN indicate greater rates of beneficial biogeochemical cycling. However, both approaches are consistent with the idea that different functions are valued in different ways. We then calculated multifunctionality as for the laboratory experiment by summing *z-*scores of each function.

### Predictors of chemodiversity and ecosystem function

Ecosystem functions and chemodiversity are often influenced by physicochemical variables^16,36^ and climate^8,9^. Therefore, during the field study, we recorded water temperature and pH using a multiprobe (HI-99171, Hanna Instruments, USA), measured water colour as the absorbance at 254nm using a spectrophotometer (Flame UV-Vis, Ocean Optics, USA), and measured chlorophyll *a* using a portable fluorometer (FluoroProbe III, bbe Moldaenke, Germany). In the lab, we measured TDN concentrations concurrent with DOC concentrations on the Shimadzu TOC-TNM-L analyser. Climate variables were the mean annual temperature (MAT) and mean annual precipitation (MAP) of each lake at a ca. 31km resolution compiled by the European Centre for Medium-Range Weather Forecasts ERA5 data product^88^. We clipped the ERA5 analysis to the area of each catchment retrieved from the Global Lake area, Climate, and Population dataset^89^ and averaged across months for all pixels in a catchment to obtain MAT and MAP. Because ecosystem function can also depend on the surrounding catchment, a net primary production (NPP) raster (500m resolution) based on MODIS satellite images^90^ was clipped to each catchment and averaged across all pixels in a catchment to derive NPP.

### Testing the effects of chemodiversity and biodiversity on ecosystem multifunctionality

We first determined which variables influenced multifunctionality in the lab experiment and field survey by fitting a linear model that only contained biodiversity and chemodiversity. We then fitted a second linear model that included an interaction between biodiversity and chemodiversity to test if any effects of chemodiversity on multifunctionality were conditional on biodiversity. For the lab experiment, chemodiversity and biodiversity were treated as factors, as we only had ≤4 levels of each variable, whereas we used them as continuous variables in the field survey. We used the same model for testing the effect of chemodiversity on multifunctionality as for testing the effect of chemodiversity on each individual function in the laboratory experiment. However, we added mean annual temperature, water colour, lake area, and chlorophyll *a* concentration to model each function in the field survey because previous studies have revealed that these variables also predict ecosystem function^23,24^. All models were fitted with R version 4.1.1^78^.

In addition to the linear models, we tested explicit hypotheses about the drivers of chemodiversity-ecosystem multifunctionality relationships using a path analysis. Here, we designed a path analysis explaining variation in multifunctionality, chemodiversity, the number of biochemical transformations, and DOM complexity. We hypothesized that variation in multifunctionality was explained by chemodiversity, as well as biodiversity, MAT, and MAP, which all predict ecosystem multifunctionality in other studies^3,10,38^. We then hypothesized that chemodiversity was explained by its complexity and potential transformation consistent with established theories for the reactivity of DOM^27^. Based on the same theories, we hypothesized that the number of times a molecule was transformed was also influenced by its complexity and all the environmental variables we measured, including biodiversity. Finally, we hypothesized that DOM complexity was influenced by terrestrial primary production that naturally produces many complex compounds that are leached into downstream waters^65^. Although microbial activity also increases complexity^13^, we expected its effect to be minimal because most DOM in lakes originates from terrestrial production^16^. We then fitted this path analysis to lake-level observations using the piecewiseSEM package^91^ in R, which translates our hypotheses into a set of linear models. After fitting the initial path analysis, we included missing paths identified by tests of directed separation as described in Fig. S13.

### Identifying the production and consumption of individual compounds

For the laboratory experiment, we first identified individual molecular formulae that were consumed or produced by microbes. We compared the relative abundance of each molecular formula at the end of the incubation experiment from all biodiversity levels against the relative abundance of the same formula in the control with a Welch’s t-test. We only considered the final time point of the control to account for compounds that could be produced or consumed simply due to abiotic processes. We considered compounds were produced and consumed if their abundance significantly increased and decreased compared to the control, respectively. We then summed the total number of produced and consumed compounds at each chemodiversity level.

For the field survey, we tested how the consumption and production of formulae was associated with each function that varied with chemodiversity across Europe (CH_4_ and CO_2_ emissions and bacterial biomass production). We correlated the relative abundance of each formula in each lake with the value of each function in the corresponding lake using Spearman rank correlations. We considered statistically significant negative and positive correlations to indicate that consumption and production of a formula, respectively, was associated with a given function.

### Extrapolating chemodiversity and ecosystem function from our field survey

To extrapolate chemodiversity and each measured function that varied with chemodiversity across Europe (CH_4_ and CO_2_ emissions and bacterial biomass production), we first fitted linear models to these variables. The models included all physio-chemical and environmental variables measured in the field survey, and we included chemodiversity and biodiversity as predictors when modelling each function. Then, we simplified the full models with stepwise variable selection guided by the Akaike information criterion (AIC). We sequentially removed the least supported variable, that is, most increased AIC when retained. If all variables reduced AIC when retained, we dropped the variable that least reduced AIC by <2 units^92^. We repeated this selection process until all variables retained in the model reduced AIC by ≥2 units. The goal of this analysis was to predict each function as accurately as possible to extrapolate across Europe rather than simply compare the importance of chemodiversity versus biodiversity as when modelling multifunctionality. Model selection indicated that chemodiversity was best explained by NPP, MAP, and chlorophyll *a* concentration (Table S3), which was identical to the variables that explained variation in chemodiversity within our path analysis (Figs 2, S13).

To extrapolate our linear models across Europe in 2019, we forced them with the same MAP and NPP rasters described above. The other predictors required in our models were water colour and chlorophyll *a* concentration (Table S3, S4), which are unavailable at a European-scale in publicly available datasets. Therefore, we predicted water colour from elevation, MAP, and lake area using a linear model identical to past global studies predicting water colour^75^. Similarly, we predicted chlorophyll *a* from MAT, elevation, MAP, and lake area using another linear model. Past global studies have used all these same variables to predict chlorophyll *a*, except that they included TP, which was also unavailable from publicly available datasets at a European-scale^23,24^. Although the models explained 25% and 29% of the variation in water colour and chlorophyll *a*, respectively, the errors introduced by using these models are not relevant for our calculations focusing on relative changes over time (see section below).

### Forecasting the future functioning of lakes

We estimated how each function that varied with chemodiversity (CH_4_ and CO_2_ emissions and bacterial biomass production) would change across Europe by 2100 solely because of changes to chemodiversity. To predict chemodiversity in 2100, we first obtained the IPSL-CM6A^60^, HadGEM3-GC31^93^, and ACCESS-ESM1.5^94^ climate forecasts for 2100. We selected these forecasts because they emphasise the relevance of terrestrial-atmosphere exchange and are frequently used in climate studies^95^. We downloaded rasters of MAT and MAP for likely (SSP2) and worst case (SSP5) scenarios for future changes in atmospheric CO_2_ concentrations for each forecast from the Copernicus Climate Data Store (https://cds.climate.copernicus.eu/). NPP predictions for each forecast were downloaded from the Earth System Grid Federation servers (https://esgf-node.llnl.gov/search/cmip6/). Because no NPP data were available for the HadGEM3-GC31 model, we used the average of the IPSL-CM6A and ACCESS-ESM1.5 models. We then input these rasters into our linear model of chemodiversity to predict chemodiversity in 2100 across Europe (Table S3). To generate the predictor of chlorophyll *a*, we again used the model described in the previous section but with MAT and MAP from the climate forecasts, as well as elevation and lake area. The chemodiversity raster was then input into the linear models of CH_4_ and CO_2_ emissions and bacterial biomass production (Table S4). All predictors other than chemodiversity were set at their 2019 values. Rather than focus on absolute values of each function, we calculated the change in each function between 2019 and 2100 when only changing chemodiversity, that is, holding all other variables constant. Because errors associated with models of water colour and chlorophyll *a* are likely to be systematic, that is, non-random, the errors should remain similar between time periods with chemodiversity being the sole difference in the comparison. Furthermore, estimates of the magnitude of changes of CH_4_ and CO_2_ for all of Europe were well in line with previous global studies^23,24^, further suggesting that errors in absolute values of water colour and chlorophyll *a* were of minor importance.

