## Supplementary material for "Chemical diversity promotes ecosystem function"

***Supplementary Note 1: Estimating ecosystem multifunctionality***

There is no consensus for how to estimate ecosystem multifunctionality^1–3^. However, two approaches predominate^2^. The first, referred to as the “average approach”, infers multifunctionality by averaging *z*-scores of all measured functions^4^. The “average approach” generates a single, continuous index of multifunctionality per site, which eases interpretation^4^. However, the resulting index is study-specific, as it depends on the functions that were averaged, thereby complicating cross-study comparisons^2^. The second approach, referred to as the “threshold approach”, evaluates how many functions exceed a specific threshold, often of a maximum value^1^. The “threshold approach” can facilitate inter-study comparisons because it is estimated in a common unit, i.e. number of functions^1,2^, but interpretation is more difficult. There is, in theory, an infinite number of thresholds that can be used to define the index and so there are as many values with this approach as there are thresholds.

In this study, we used the average approach for its simple interpretation. Nevertheless, to allow for comparison with other studies, we also computed multifunctionality using the “threshold approach”. We used the *z*-scores of all measured variables and assessed how many functions were higher than 1% to 99% (in 1% intervals) of their respective maximal value. For each function, we excluded the five largest values to avoid bias introduced by potential outliers, as recommended for this approach ^2^. For the same reason, we averaged the five largest values to estimate the maximum values a function can reach. We then fitted a linear model to predict the threshold multifunctionality from chemodiversity and extracted the estimated slope. As in the Main Text (i.e. Fig. 1), we accounted for variation in multifunctionality simply due to MAT. We also considered clustering and weighting the functions before using the “threshold approach” as recommended where two or more functions measure similar processes^2^. However, the five measured functions corresponded to distinct biogeochemical processes, despite all being related to carbon cycling, and so we did not cluster them together.

We found that multifunctionality increased with chemodiversity at high but not low thresholds. Specifically, chemodiversity had a positive effect on the number of functions exceeding all thresholds above a maximal value for the functions of 79% (Fig. S19). These results suggest that to sustain multifunctionality at a high level (>79% of maximal values), novel molecular formulas are required. The present study is the first to evaluate the effects of chemodiversity on multifunctionality and, therefore, we cannot compare our data with others. However, we can relate our findings to biodiversity studies. Specifically, in comparison to a synthesis of 47 studies^5^, chemodiversity sustained multifunctionality at a higher possible threshold (99%) compared with 94% in the latter. Another metric used to compare the effect of diversity on multifunctionality is the threshold at which diversity has the maximal effect on multifunctionality T_mde_. A larger T_mde_ indicates a stronger effect of diversity on multifunctionality^1^. In our study, T_mde_ for chemodiversity was 91%, as compared with 63% for biodiversity and <70% for the effect of biodiversity on multifunctionality estimated from a similar number of functions in the cross-ecosystem synthesis^5^. Overall, our results confirm that chemodiversity has a stronger effect on multifunctionality than biodiversity and that increase chemodiversity can sustain higher levels of multifunctionality than biodiversity.

***Supplementary Note 2: Intrinsic and emergent recalcitrance hypothesis***

The emergent recalcitrance hypothesis states that chemodiversity at large spatial and temporal scales is mainly controlled by the properties of microbes that transform DOM and the environmental conditions that influence these interactions^6,7^. At higher latitudes, different microbes and different environmental conditions, such as lower mean annual temperatures (MAT) and lower macronutrient availability, could slow microbial use of DOM and cause more molecular formulae to persist. Concurrently, the intrinsic recalcitrance hypothesis states that chemodiversity depends on the inherent molecular properties of DOM that determine the extent to which microbes transform individual substrates into a diversity of compounds^6,7^. Terrestrial vegetation, which is often the main source of lake DOM^8^, tends to invest more in protecting biomass, such as with complex polyphenolics^9^, as plants face a higher cost to replace biomass, e.g., as MAT declines^10^. These more complex compounds have properties including a greater degree of unsaturation, lower ratio of hydrogen to carbon atoms, and higher oxidation state of carbon, all of which make them more difficult to be biochemically transformed by microbes or photochemical processes^11^. Chemodiversity therefore increased at higher latitudes (Fig. S11) where diverse and complex compounds that are more difficult for microbes to transform were produced. In the present study, we utilized the concept of intrinsic and emergent recalcitrance to formulate the hypothesis that knowledge of the origin of chemodiversity should help us understand how chemodiversity influences ecosystem multifunctionality.

***Supplementary References***

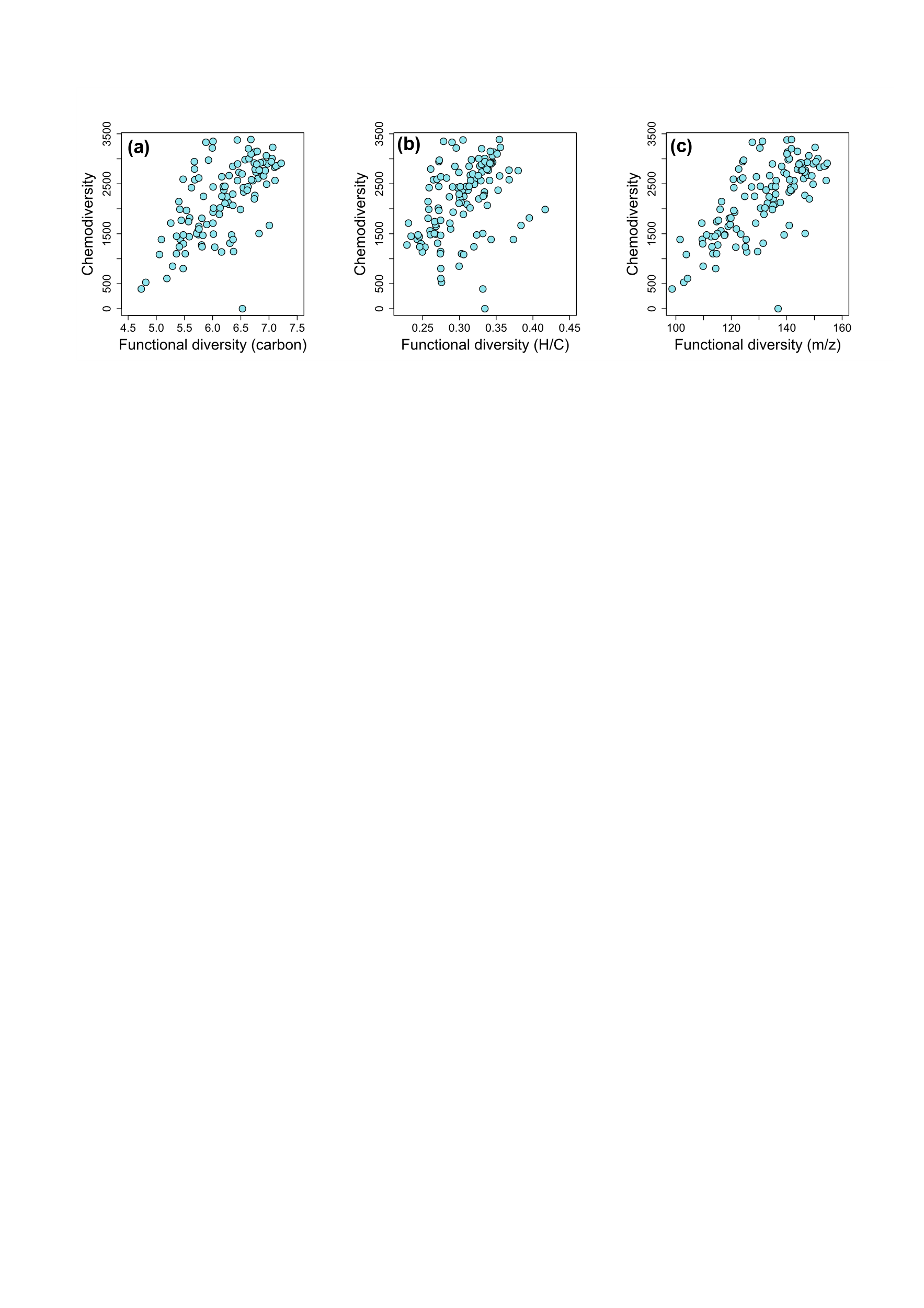

**Figure S1: Chemodiversity correlates with the functional diversity of dissolved organic matter**. For each lake, we computed three measures of functional diversity based on the chemical properties of their molecular formulae. The measures estimated the diversity in the **(a)** sizes of compounds, as inferred from the number of carbon atoms they contained, which reflects their reactivity^12^; **(b)** the bioavailability of molecular formulae, whereby formulae with a greater ratio of hydrogen to carbon elements (H/C) are considered more easily degraded^11^, and **(c)** the molecular masses (m/z) of compounds, which reflects the origins and degree of transformation of molecules^11–13^. We used Rao’s quadratic entropy to calculate each functional diversity measure from the presence and relative abundance of each molecule in each lake following Mentges *et al*^14^. Each functional diversity index was closely correlated with chemodiversity with Spearman rank correlation coefficients of 0.63, 0.46, and 0.69 for **(a)**, **(b)**, and **(c)**, respectively, with p-values <0.001 for all.

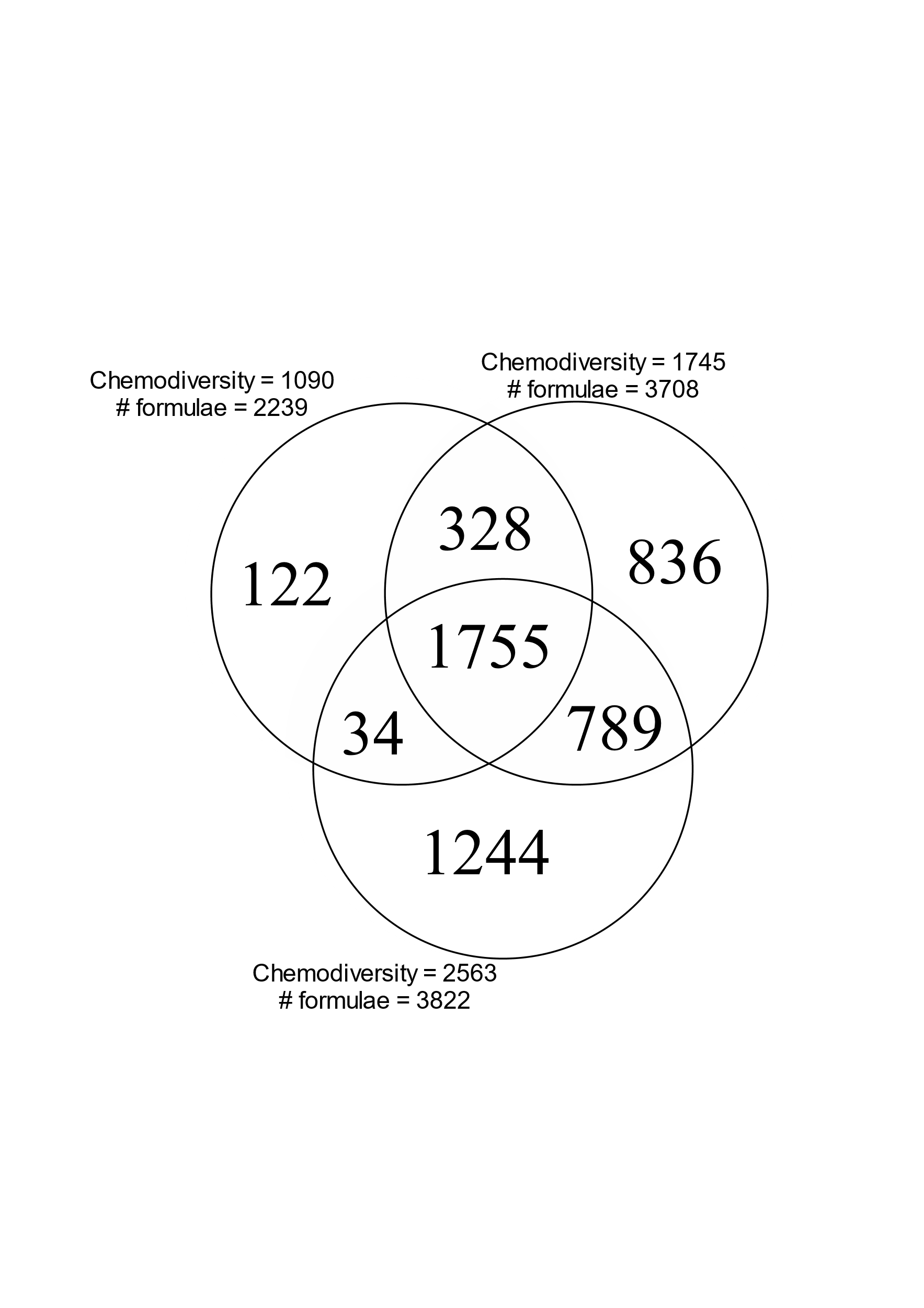

**Figure S2: The three experimental DOM sources were compositionally nested**. In other words, increasing chemodiversity did not entirely change molecular composition but added new compounds onto those present in treatments with lower chemodiversity. Circles indicate the number of molecular formulae shared between the different chemodiversity sources and correspond to the low chemodiversity (i.e. exponential for the Shannon diversity index = 1090) Arctic river, the medium chemodiversity (i.e. 1745) boreal lake, and the high chemodiversity (2563) temperate wetland. The exponential of the Shannon diversity index was used to estimate chemodiversity and is equal to the effective number of molecular formulae. This value is less than the observed number of molecular formulae because it corrects the diversity estimate for variation in evenness in the relative abundance of each formula.

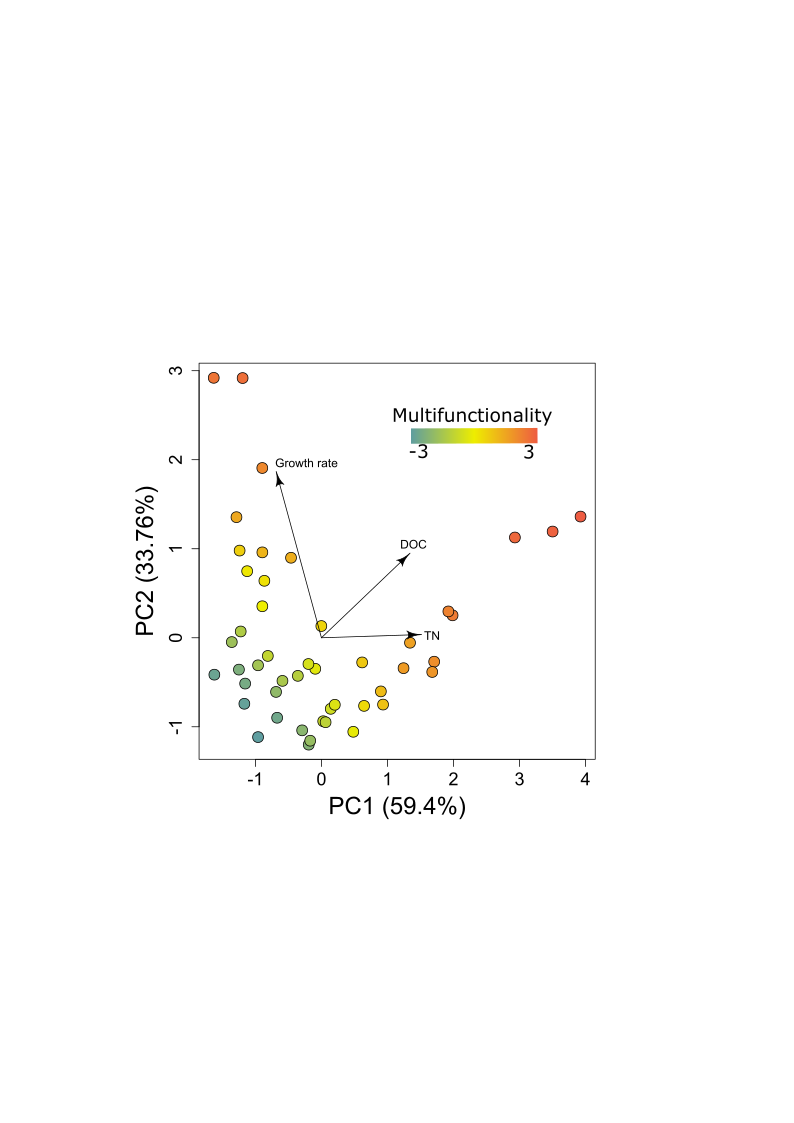

**Figure S3: Ecosystem functions measured in the laboratory experiment were largely orthogonal to each other.** We used a principal component analysis to visualize the associations among three ecosystem functions measuring the change in each of cell abundances, dissolved organic carbon (DOC) concentrations, and total nitrogen (TN) concentrations over a 7-day period. Each of the N=36 dots correspond to one Erlenmeyer flask. The colour gradient indicates multifunctionality, with warmer colours indicating higher multifunctionality. The variation explained by each principal component (PC) axis is indicated along the axes. In our experiment, DOC and TN correlated positively (ρ=0.55, p<0.001) and TN and growth rate correlated negatively (ρ=-0.41, p=0.005).

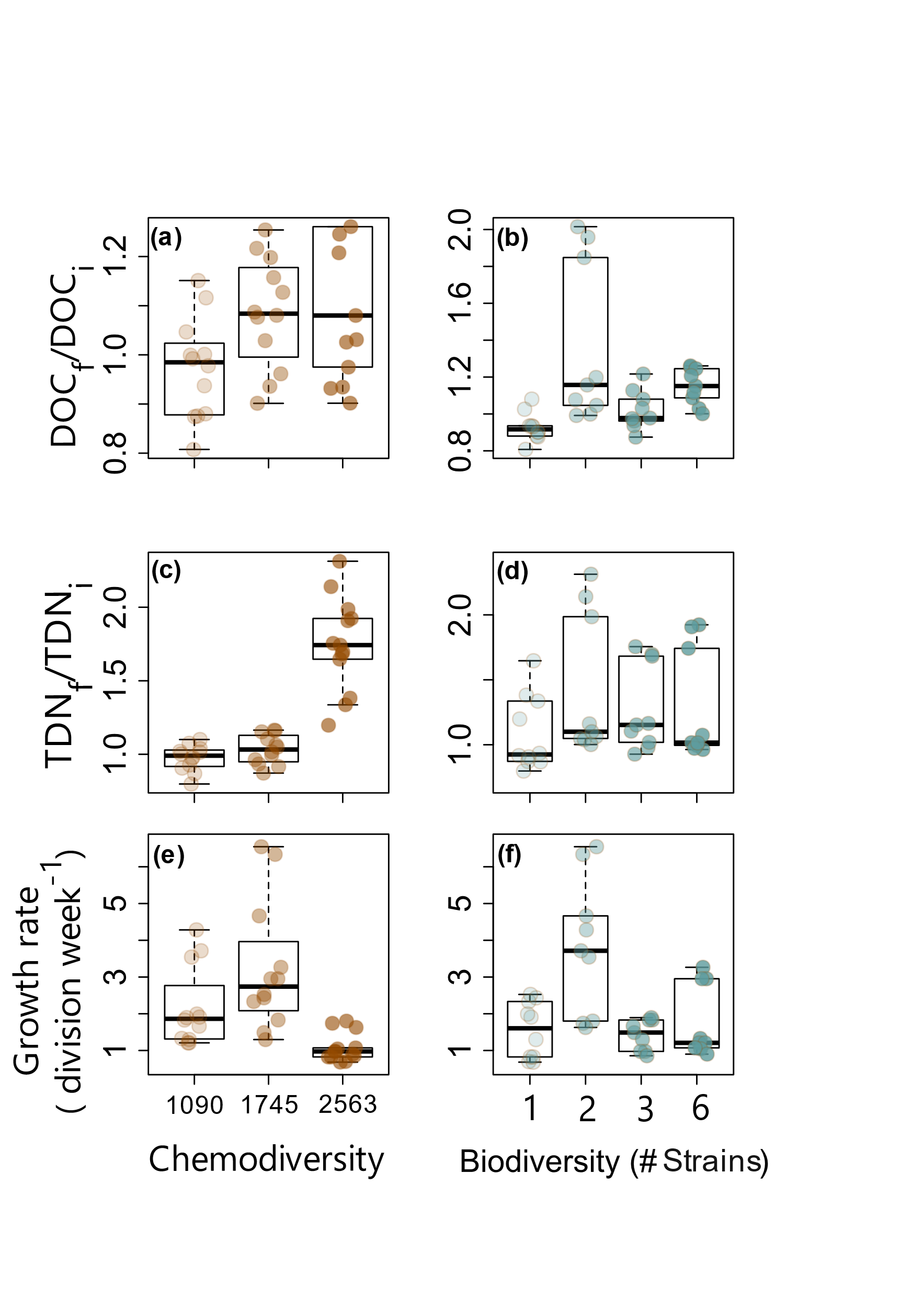

**Figure S4: Increasing chemodiversity is more often associated with higher ecosystem function than biodiversity in a laboratory experiment.** We tested how each of dissolved organic carbon (DOC) consumption, total dissolved nitrogen (TDN) consumption, and growth rate (cell divisions per week) varied with chemodiversity source and biodiversity using linear models. Consumption was measured as the ratio of final (f), i.e. after 7 days, versus initial (i) measurements. Increasing chemodiversity increased both DOC (t=2.9, df=35, p=0.006) and TDN (t=8.2, df=35, p<0.001) consumption and decreased growth rate (t=-2.2, df=35, p=0.046). The observed growth rates were relatively low because we exposed microbial communities to natural organic matter without additional nutrients rather than a growth media and the decline with increasing chemodiversity can be attributed to the corresponding increase in the proportion of aromatic compounds (Table S1). Although more compounds are used by the bacterial communities at higher chemodiversity (Fig. S5), the degradation of those compounds does not provide as much energy to microbes as the degradation of more aliphatic compounds at lower chemodiversity (Table S1). Biodiversity had no effect overall on DOC (t=0.7, df=35, p=0.451) and TDN (t=0.6, df=35, p=0.558) consumption, or growth rate (t=-0.8, df=35, p=0.433). However, the two-species community had higher levels of all three functions compared to when these species were included in the six-species community (t=3.1, p=0.018, t=6.7, p<0.001, and t=4.9, p=0.002, for DOC, TDN, and growth rate, respectively). Darker colours indicate increasing chemodiversity (brown) and biodiversity (blue). Boxes represent the median, 25^th^, and 75^th^ percentiles, and whiskers extend 1.5-times the interquartile range. Points are individual observations (total n = 36).

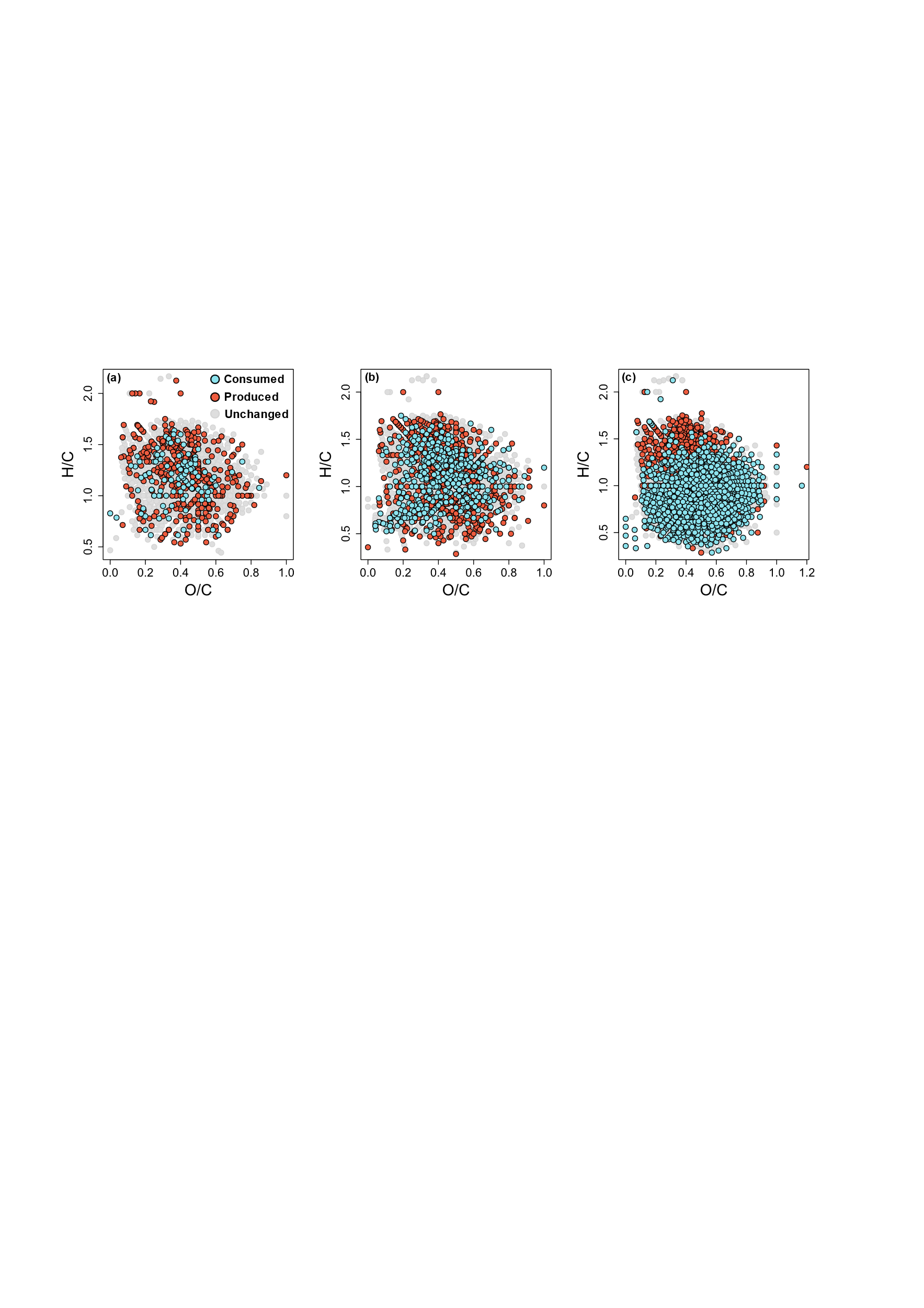

**Figure S5: Increasing chemodiversity increases the number of compounds transformed by microbes in the laboratory experiment.** We identified molecular formulae (points) that decreased (blue), increased (red), or were unchanged (grey) in relative intensity after 7 days compared to the control incubations using t-tests (p<0.05). Chemodiversity increased the number of compounds consumed with 114, 432, and 1838 compounds consumed in the sources with an estimated chemodiversity of **(a)** 1090, **(b)** 1745, and **(c)** 2564 molecular formulae, respectively. Chemodiversity did not affect the number of compounds produced, with 365, 689, and 486 compounds produced in the sources with a chemodiversity of **(a)** 1090, **(b)** 1745, and **(c)** 2564, respectively. Compounds produced and consumed during the experiment occupied all regions of the van Krevelen diagram, suggesting that microbes consumed a diverse range of compounds. Additionally, we found that 9%, 10%, and 14% of produced compounds contained nitrogen, whereas 9%, 6%, and 2% of consumed compounds contained nitrogen for the sources with a chemodiversity of 1090, 1745, and 2564, respectively. Therefore, nitrogen was incorporated into the DOM pool and increasingly so at higher chemodiversity, thereby explaining why it increased in absolute concentration (Fig. S4c,d).

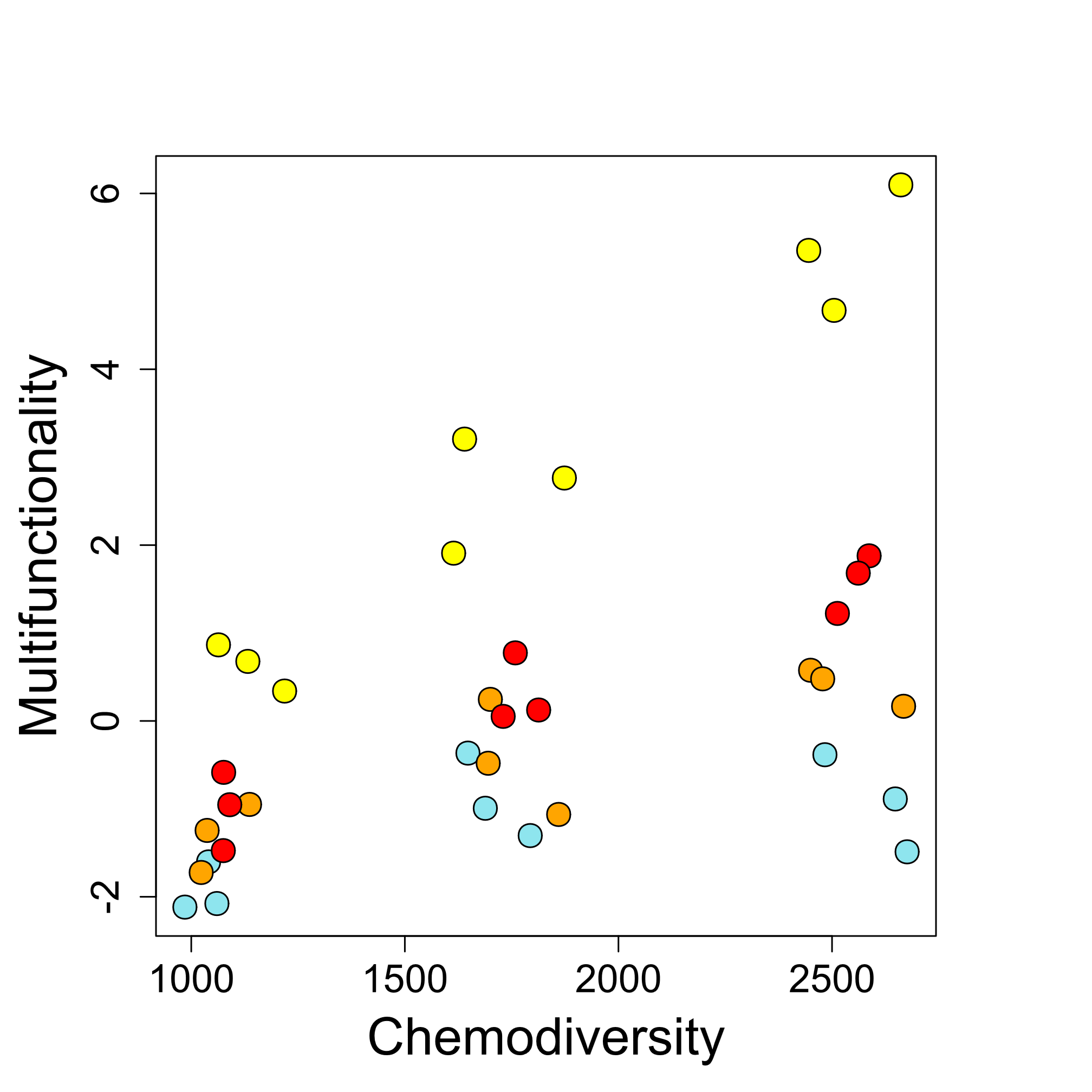

**Figure S6: Higher biodiversity increases ecosystem multifunctionality only at high chemodiversity in a lab experiment.** We tested the interaction between levels of chemodiversity and biodiversity with an ANOVA followed by a Tukey’s post-hoc test. Biodiversity was a factor with four levels: one species (blue), two species (yellow), three species (orange), and six species (red). Chemodiversity had three levels (1090, 1745, and 2564 molecular formulae) and is jittered on the *x*-axis for visualization only. The ANOVA revealed a statistically significant interaction between chemodiversity and biodiversity (F=9.9, df=6, p<0.001). At the lowest and intermediate chemodiversity levels, the two species community had a higher multifunctionality compared to the one, three, and six species communities (Tukey’s test, p<0.001). Multifunctionality observed for the one, three, and six species communities were not statistically different (p>0.100) at the lowest and intermediate chemodiversity. At the highest chemodiversity level, multifunctionality observed for the two species community remained higher than the one, three, and six species communities (p<0.001), and the multifunctionality of the three and six species communities were higher than the multifunctionality of the one species community (p=0.044 and p<0.001, respectively). We also tested if increasing chemodiversity affected the multifunctionality performed at each biodiversity level. We fitted a separate linear model between multifunctionality and chemodiversity for each biodiversity level. We found no evidence that increasing chemodiversity increased the multifunctionality performed by the one species community (p>0.05 for all pairwise comparisons). However, the multifunctionality performed by the two and six species communities systematically increased as chemodiversity increased over the three levels in our experiment (p<0.001). For the three species communities, multifunctionality increased as chemodiversity increased from 1090 to 1745 molecular formulae (p=0.020), but did not increase when chemodiversity increased from 1745 to 2564 (p=0.140).

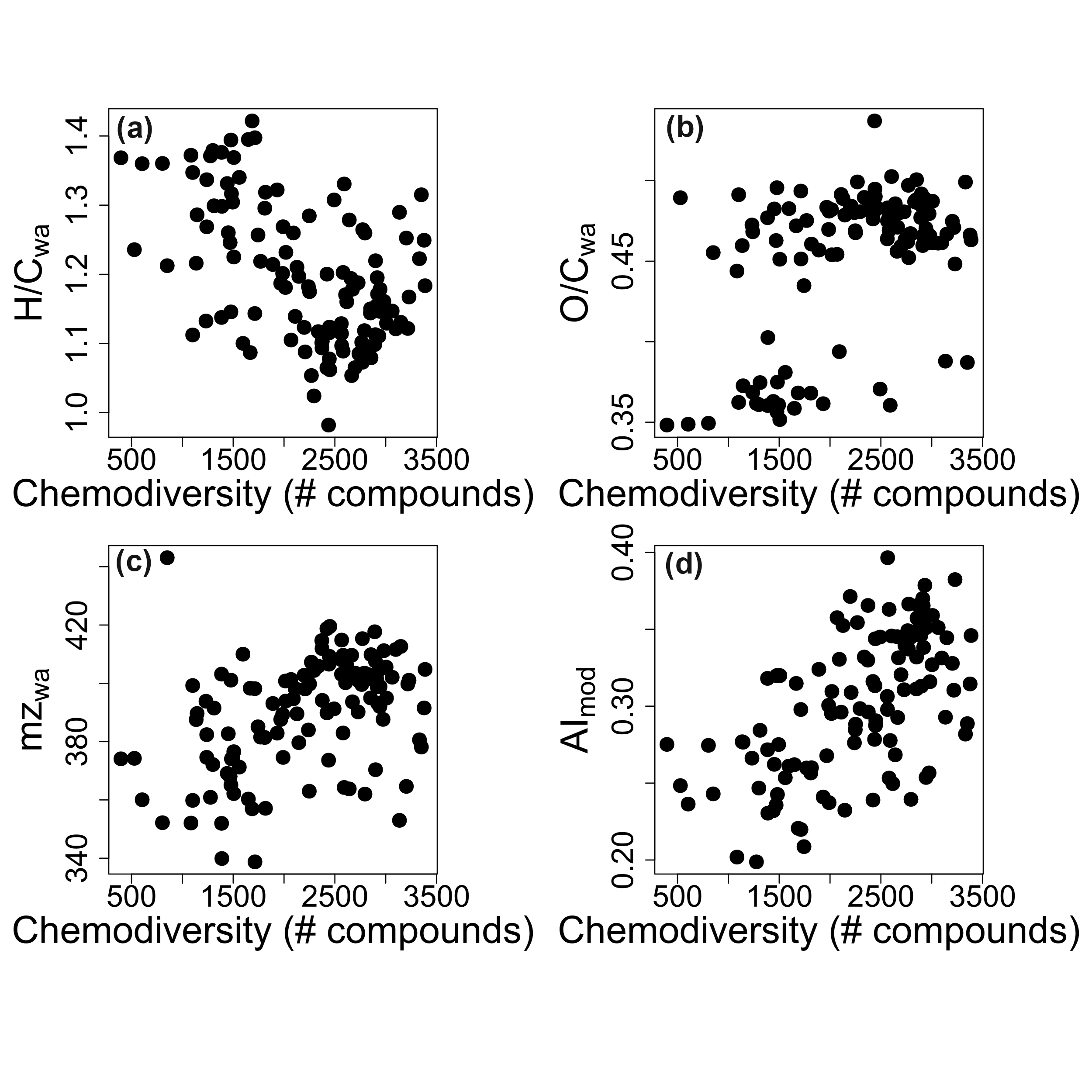

**Figure S7: The chemical characteristics of DOM vary with chemodiversity in lakes.** Chemodiversity correlated **(a)** negatively with the intensity-weighted average (wa) H:C ratio (ρ=-0.44, p<0.001), and positively with the **(b)** intensity-weighted average O:C ratio (ρ=0.34, p<0.001), **(c)** intensity-weighted average molecular mass (mz) (ρ=0.43, p<0.001), and **(d)** aromaticity index (AImod) (ρ=0.57, p<0.001). Points are individual lakes (N=101). Patterns with increasing chemodiversity were similar to those observed in the lab experiment (Table S1).

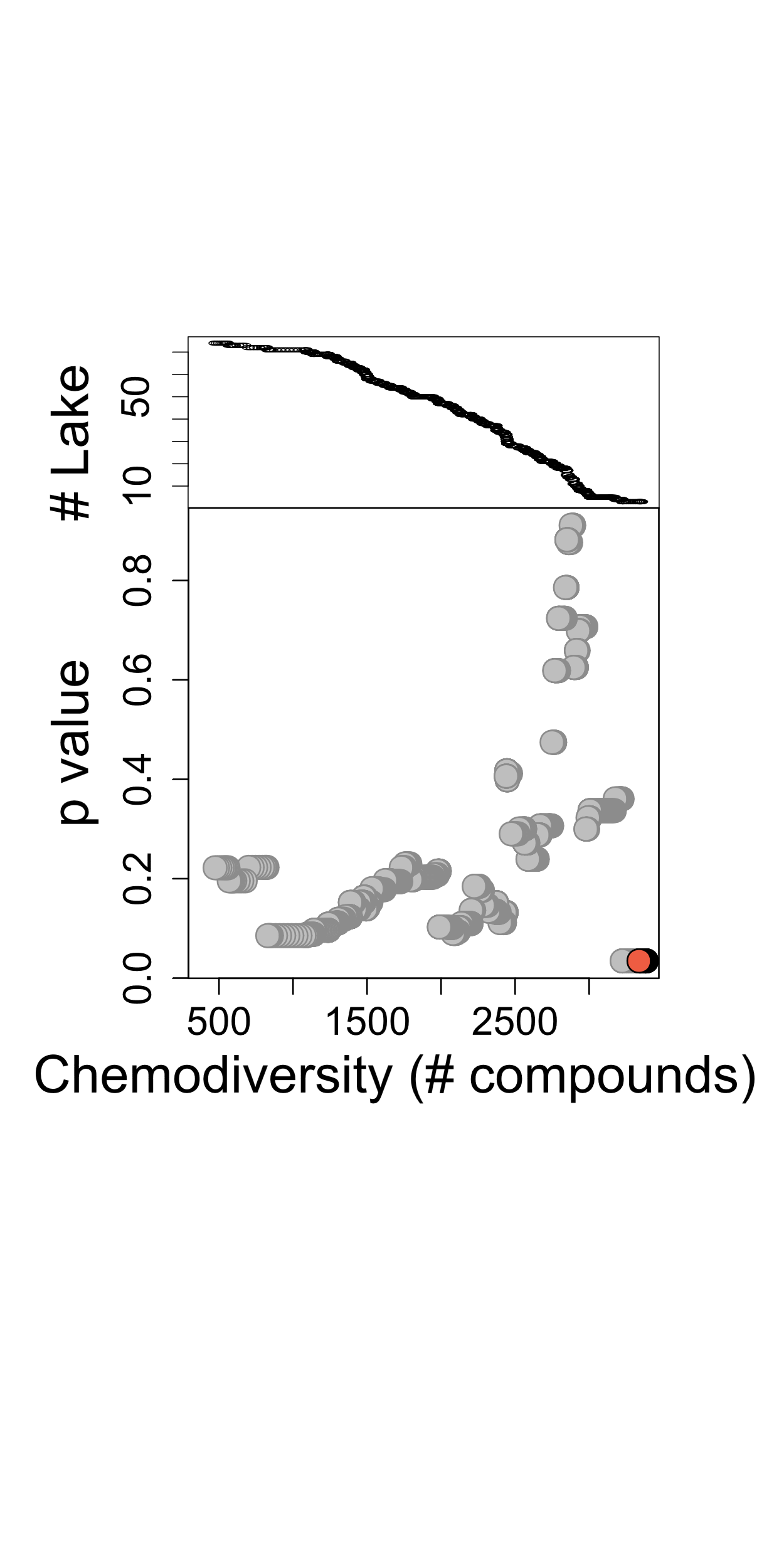

**Figure S8: Effects of biodiversity on multifunctionality increase as chemodiversity increases across lakes.** We modelled the effect of biodiversity on ecosystem multifunctionality for lakes with a chemodiversity exceeding a given threshold using linear models. The chemodiversity threshold was defined based on a quantile ranging from 0.001 to 0.999. Then, we fitted a linear model to predict multifunctionality from biodiversity using all lakes that had chemodiversity higher than the threshold. We repeated the process for each of 1000 chemodiversity thresholds. Each dot represents the p value of the effect of biodiversity on ecosystem multifunctionality at each chemodiversity threshold shown on the x-axis. Red dots indicate models with a p value lower than 0.05. The upper panel represents the number of lakes used to fit each model.

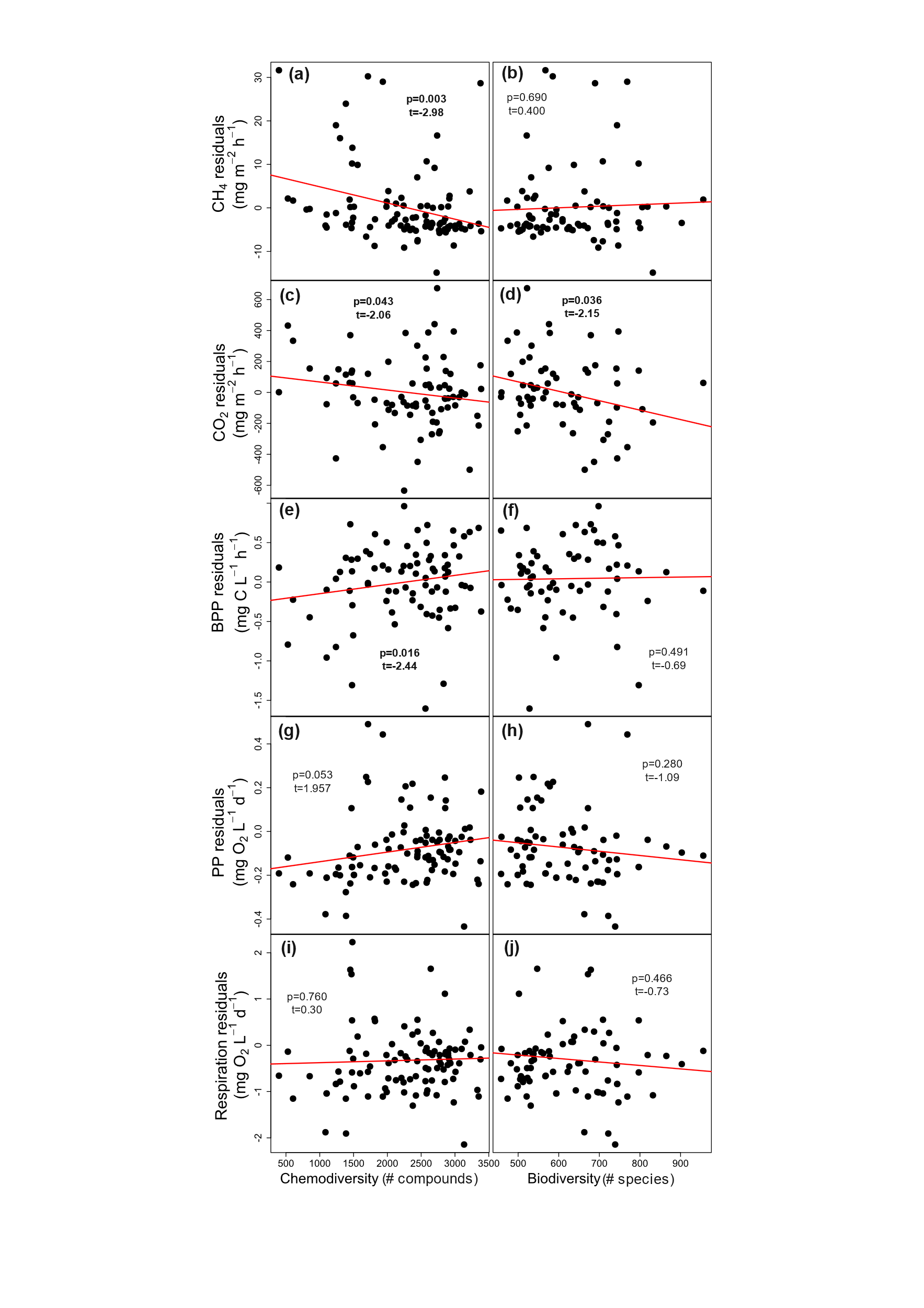

**Figure S9: Chemodiversity promoted three ecosystem functions in the field survey.** Points are lake-level (N=101) partial residuals of **(a, b)** CH_4_ and **(c, d)** CO_2_ emissions **(e, f)** bacterial protein production (BPP), **(g, h)** in-lake primary production (PP) and **(i, j)** respiration. Residuals were estimated from linear models accounting for variation in each function because of mean annual temperature, lake area, lake depth, and water color independent of chemodiversity. Lines are mean effect of chemodiversity. We modelled untransformed values of CH_4_ and CO_2_ despite reversing their signs to calculate multifunctionality. Consequently, we observed that higher chemodiversity caused lower **(a)** CH_4_ and **(c)** CO_2_ emissions, which we consider higher ecosystem multifunctionality. By contrast, biodiversity was only associated **(d)** CO_2_ emissions.

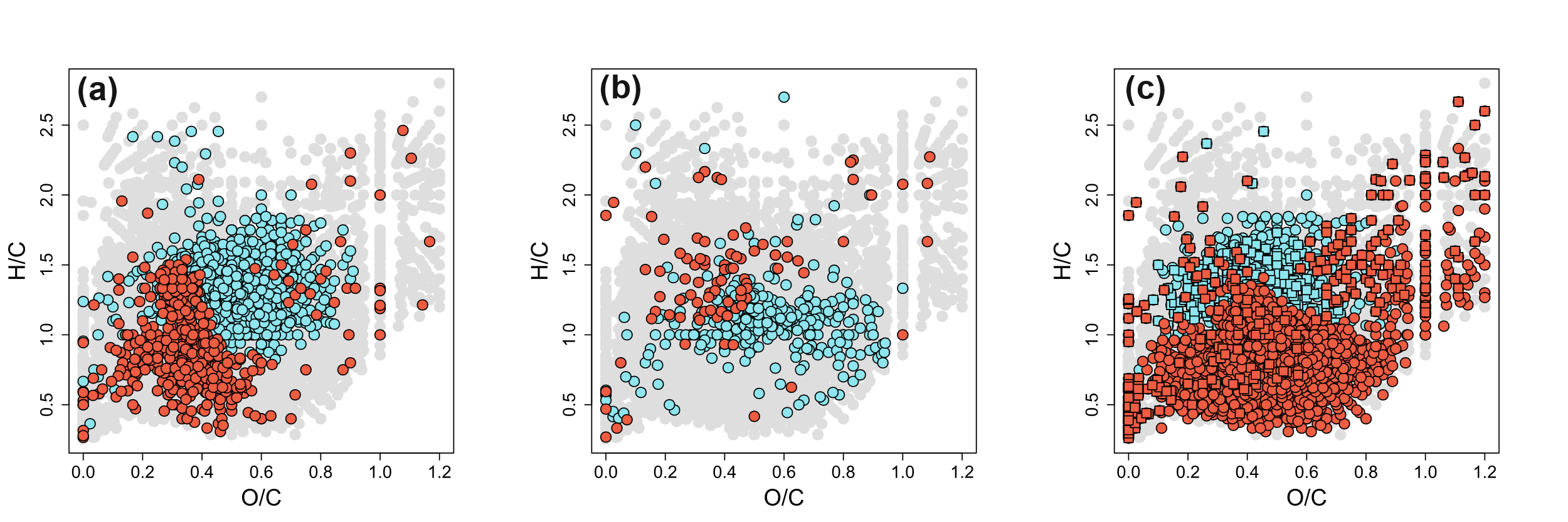

**Figure S10: Ecosystem functions related to chemodiversity associate with a specific molecular composition of DOM.** We identified compounds whose relative abundance in each lake had a statistically significant (p<0.05) Spearman correlation with **(a)** bacterial protein production, **(b)** CO_2_ emissions, and **(c)** CH_4_ emissions. Each dot represents one molecular formula (N=8456) colored according to whether the correlation was significantly positive (blue, indicative of production), negative (red, indicative of consumption), or not statistically significant (grey). In **(c)**, squares and triangles indicate formulae containing N and S, respectively. We found that the H/C of produced compounds was higher than that of consumed compounds for both bacterial protein production (t=20.7, df=685, p<0.001) and CH_4_ emissions (t=2.3, df=292, p=0.021), but lower than that of consumed compounds for CO_2_ emissions (t=-6.7, df=115, p<0.001).

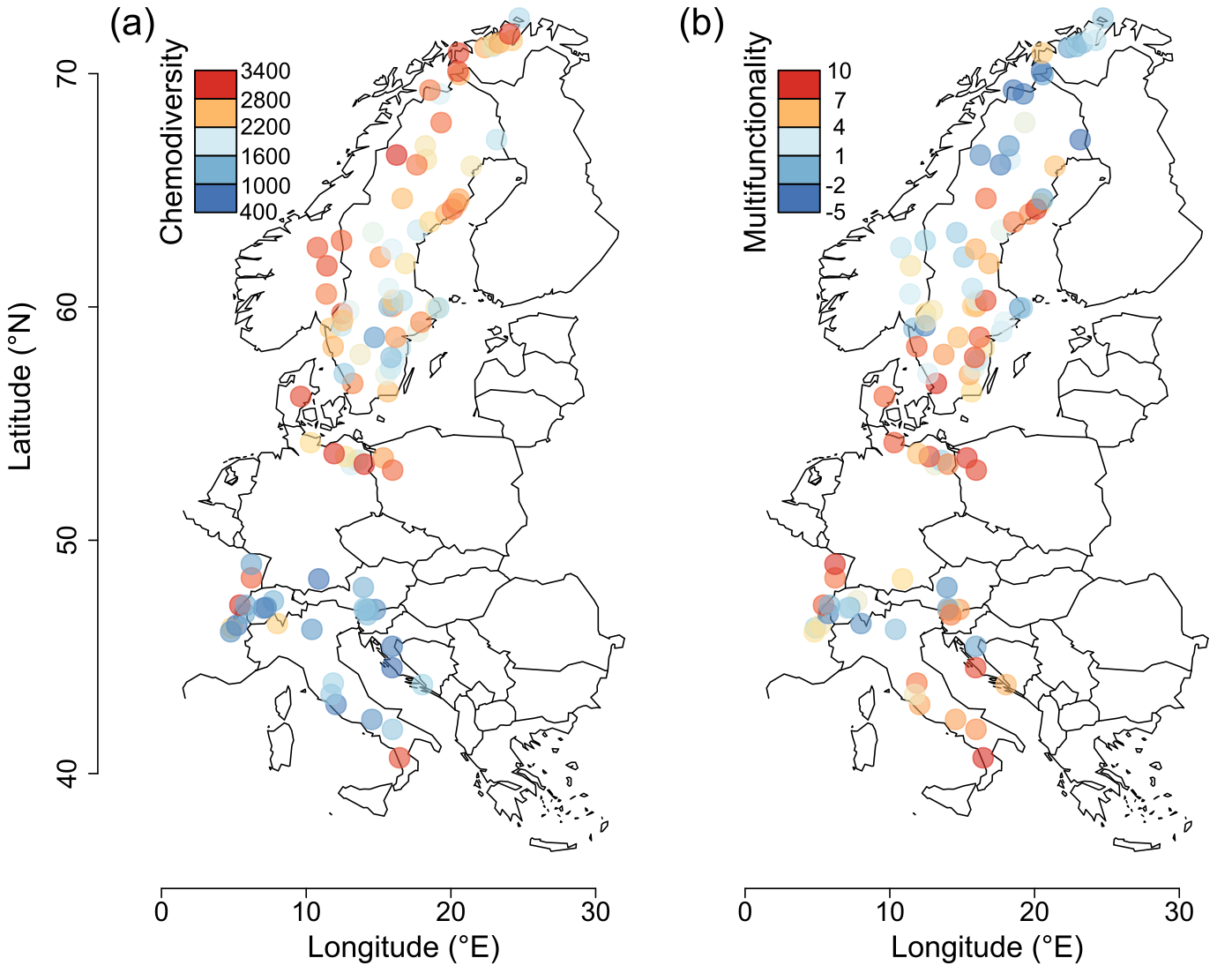

**Figure S11: Chemodiversity and multifunctionality co-vary with latitude across European lakes. (a)** Chemodiversity was calculated as the exponential of the Shannon index and so represents the effective number of different molecular formulas in dissolved organic matter. **(b)** Multifunctionality was the average z-score of five functions associated with carbon cycling and ecosystem metabolism and is dimensionless. N=101. Spearman rank correlation between chemodiversity and multifunctionality: ρ=0.24, p=0.022.

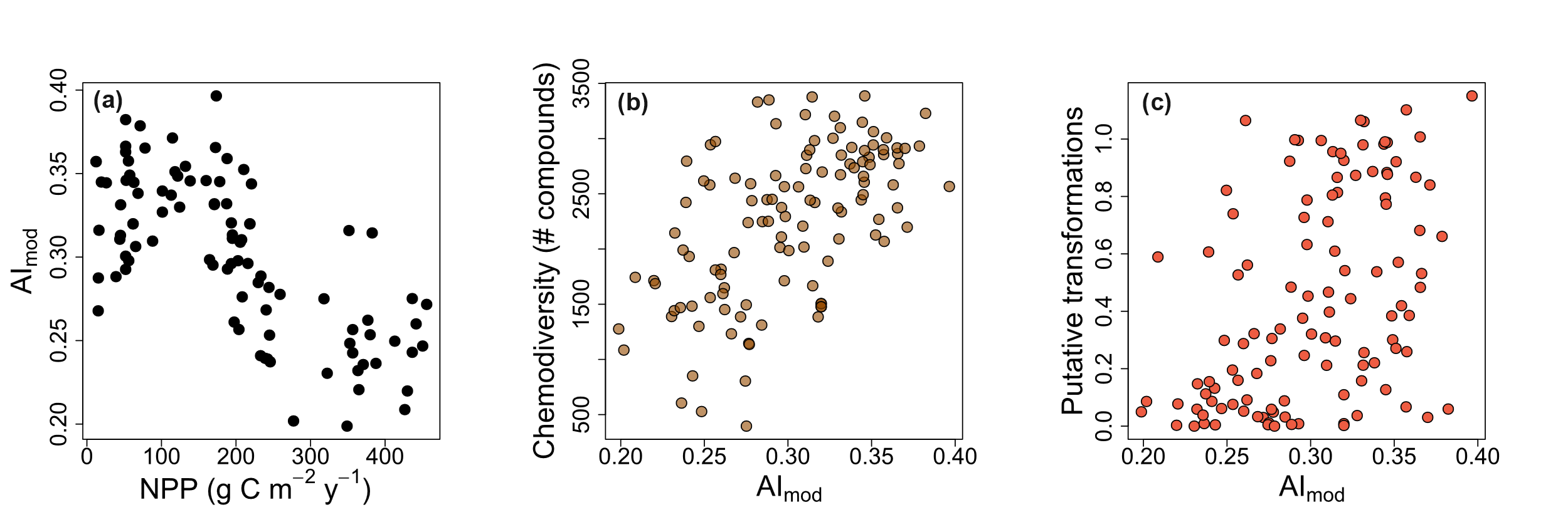

**Figure S12: DOM complexity associated with terrestrial primary production correlates with chemodiversity and the transformation of DOM. (a)** The aromaticity index (AI_mod_) reflecting DOM complexity decreased with terrestrial net primary production (NPP), consistent with the expectation that slower growing plants produce more complex compounds. **(b)** Chemodiversity increased with more complex compounds as measured by AI_mod_. **(c)** The number of putative transformations increased with complexity. The number of transformations was equal to the average number of times a molecular formula could be transformed in each lake (see Methods). Each dot represents one study lake. N=101.

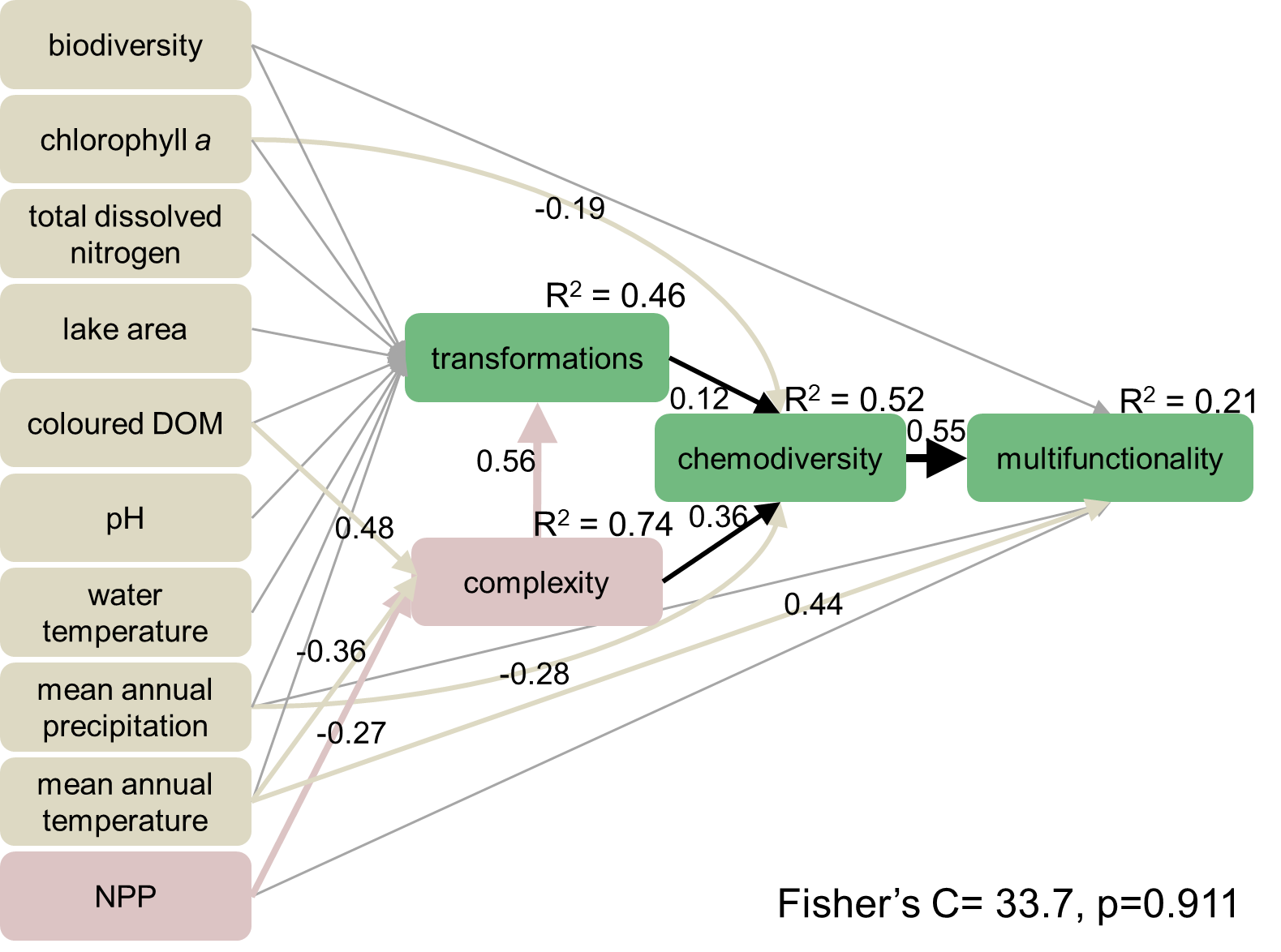

**Figure S13: Path analysis testing the drivers of the chemodiversity and ecosystem multifunctionality relationship.** We initially hypothesised that the effect of chemodiversity on multifunctionality was influenced by the molecular complexity of DOM, as measured with the aromaticity index, which itself depended on terrestrial net primary production (NPP). We also hypothesized that chemodiversity varied with the number of putative biochemical transformations per molecular formulae, which was also influenced by complexity. We included links between multifunctionality and mean annual temperature, mean annual precipitation, and biodiversity, as those variables are known to explain variation in multifunctionality^4,16,17^. However, the model structure poorly described the data due to missing paths (Fisher’s C=129.2, p<0.001), so we added paths between environmental variables, complexity, and chemodiversity based on tests of directed separation. Arrows point at modelled variables with arrows originating at predictors. Coloured arrows indicate statistically significant associations with numbers adjacent to arrows indicating mean standardised effect size and direction.
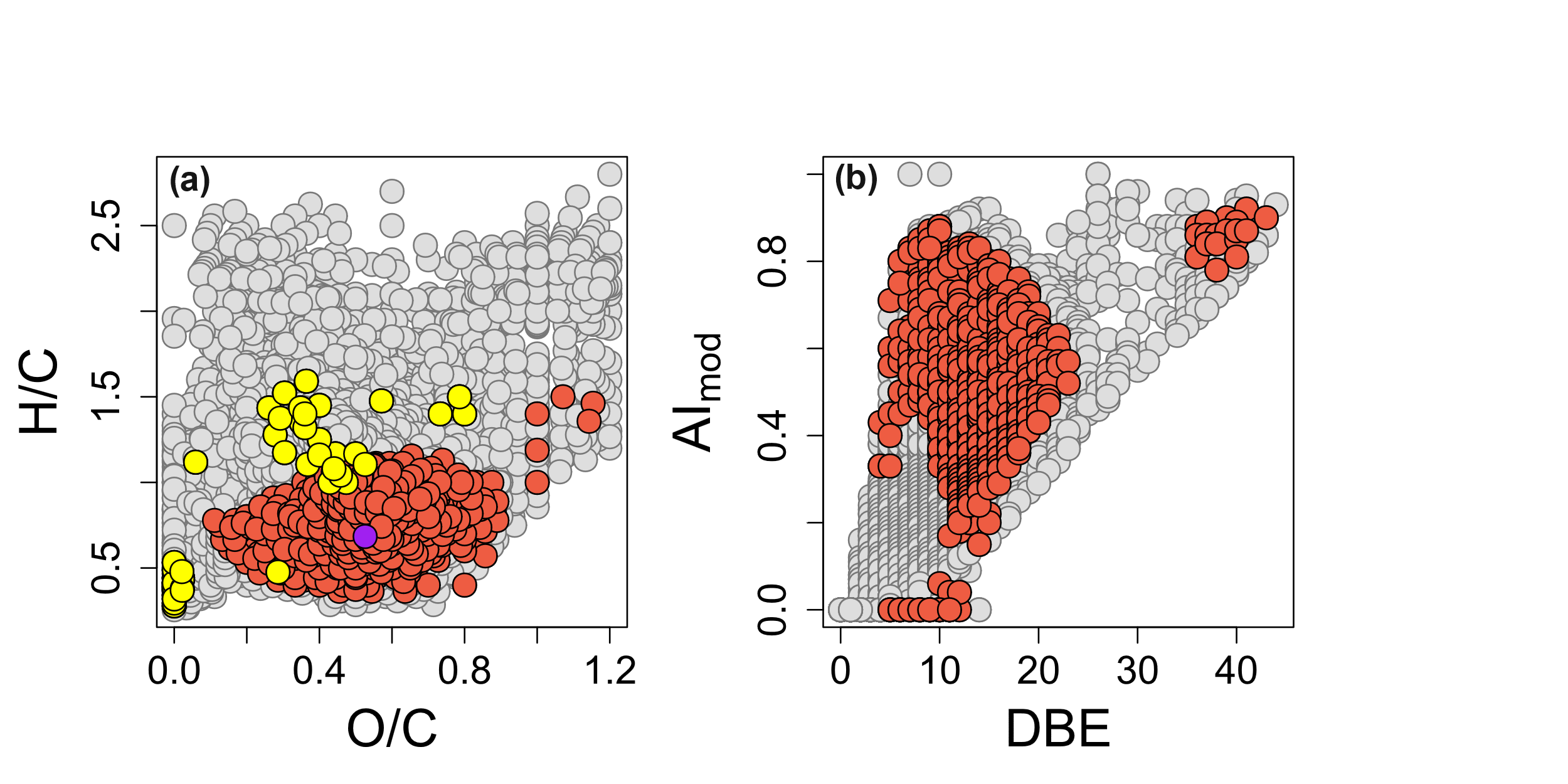

**Figure S14: Low terrestrial primary production correlates with complex compounds.** We correlated the relative abundance of each molecular formula in each lake with net primary production (NPP) in the surrounding catchment. Coloured points are formulae within the top 10% of the most negative correlations with NPP (N=846). These formulae had **(a)** a lower H/C ratio than the other formulae in our field survey (grey points), with 1 compound containing nitrogen (purple circle) and 45 compounds containing both sulphur and nitrogen (yellow circles). **(b)** Compounds associated with lower NPP also had, on average, a higher aromaticity index (AI_mod_; mean ± standard error = 0.52 ± 0.17) and double bound equivalent (DBE; mean = 14 ± 5) than other compounds identified in our field survey (AI_mod_= 0.29 ± 0.23; DBE = 10 ± 6).

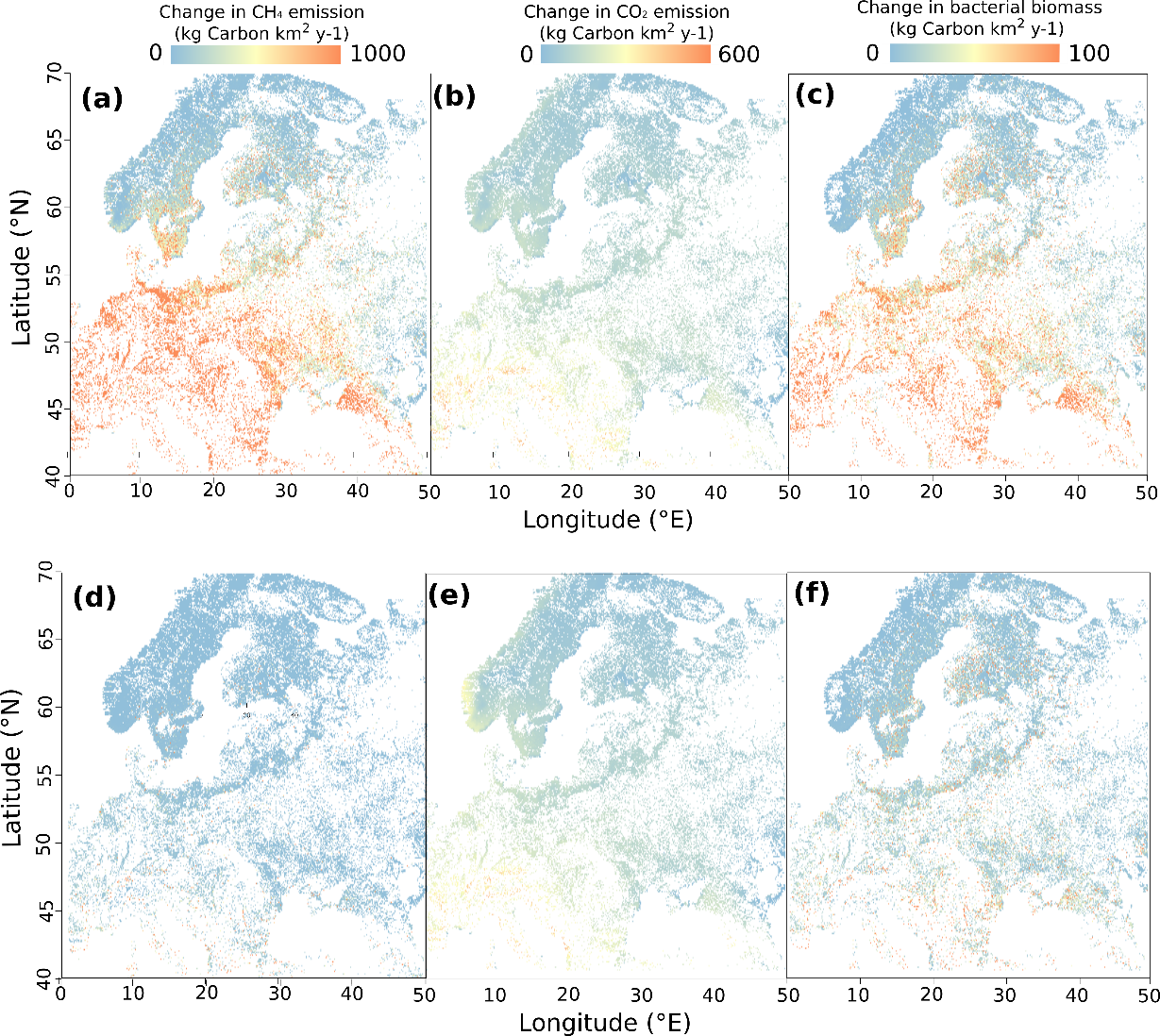

**Figure S15: Error associated with changes in ecosystem functions predicted by 2100 solely because of changes in chemodiversity.** We calculated **(a, b, c)** upper and **(d, e, f)** lower 95% confidence intervals for future changes in **(a, d)** CH_4_, **(b, e)** CO_2_, and **(c, f)** bacterial biomass production. Colours indicate the intensity of changes between 2019 and 2100. Points are individual lakes (N=175,500).

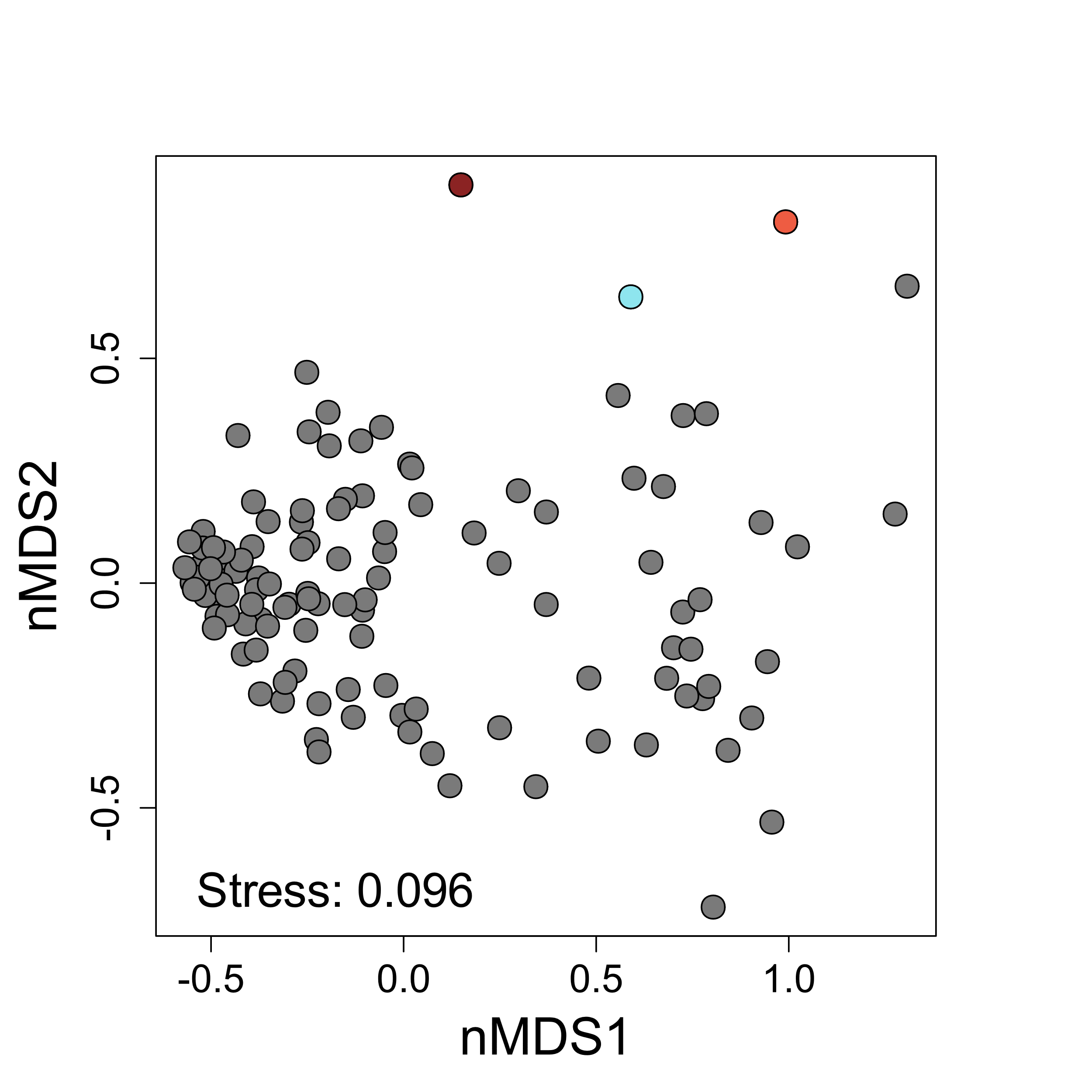

**Figure S16: The DOM sources used in the experiment were similar to European lakes.** We visualised the distance between each sampled lakes or DOM sources using non-metric multidimensional scaling (nMDS) with Bray-Curtis dissimilarities. Each grey dots represents a lake our European field survey, brown = temperate marsh, blue = boreal lake, red = Arctic river.

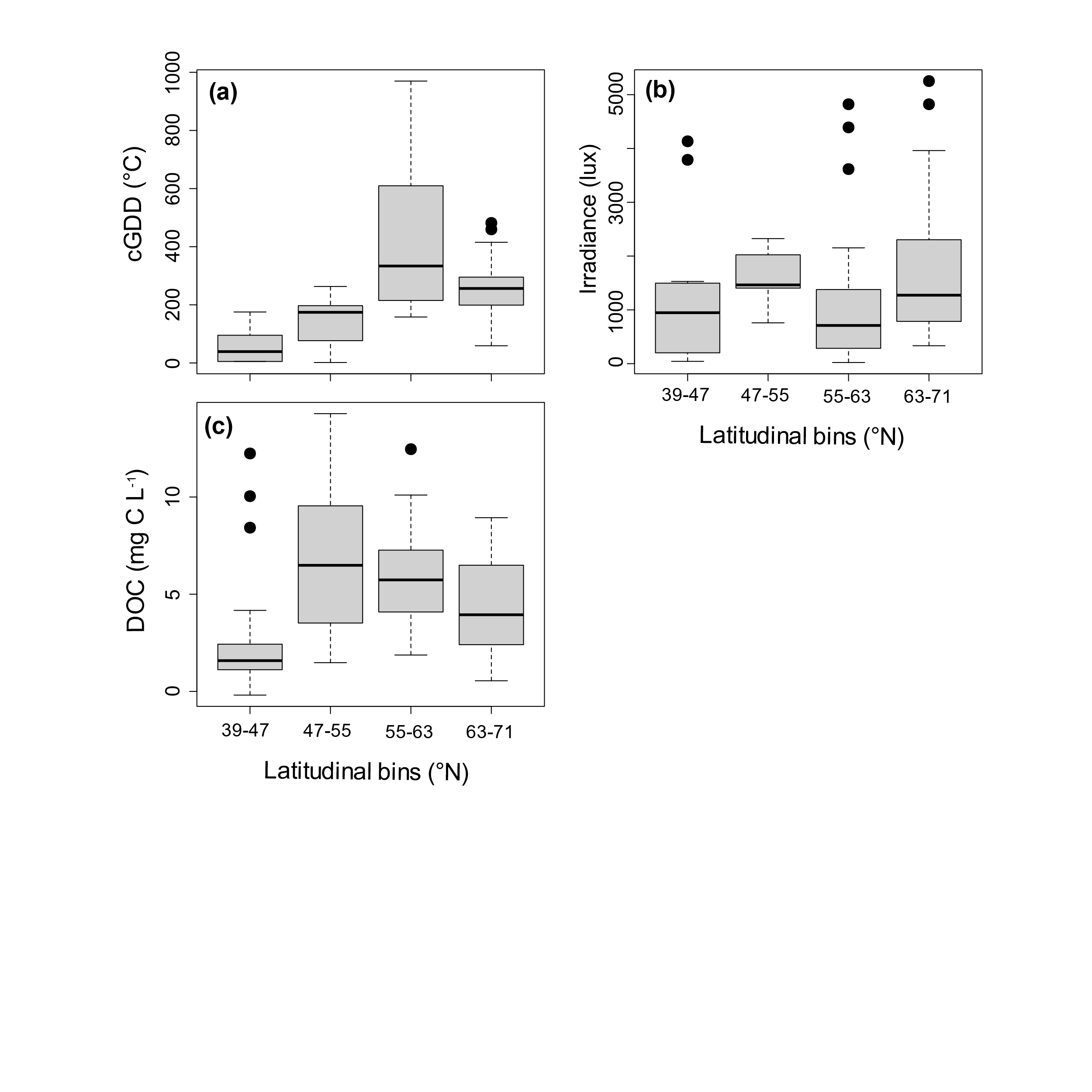

**Figure S17:** **Environmental factors did not vary monotonically with latitude.** Lakes were sampled at similar periods of **(a)** annual heat accumulation, that is, cumulative growing degree days (cGDD) and with **(b)** similar light environment and **(c)** dissolved organic carbon (DOC) concentration. Before sampling, lakes were grouped into four latitudinal bins to standardise site selection (see Methods). The 39-47, 47-55, 55-63, and 63-71 latitudinal bins contained 27, 17, 61, and 46 lakes, respectively. The horizontal line is equal to the median with boxes spanning the 25^th^ and 75^th^ percentiles. Whiskers extend 1.5 times the interquartile range and points indicate observations outside this range.

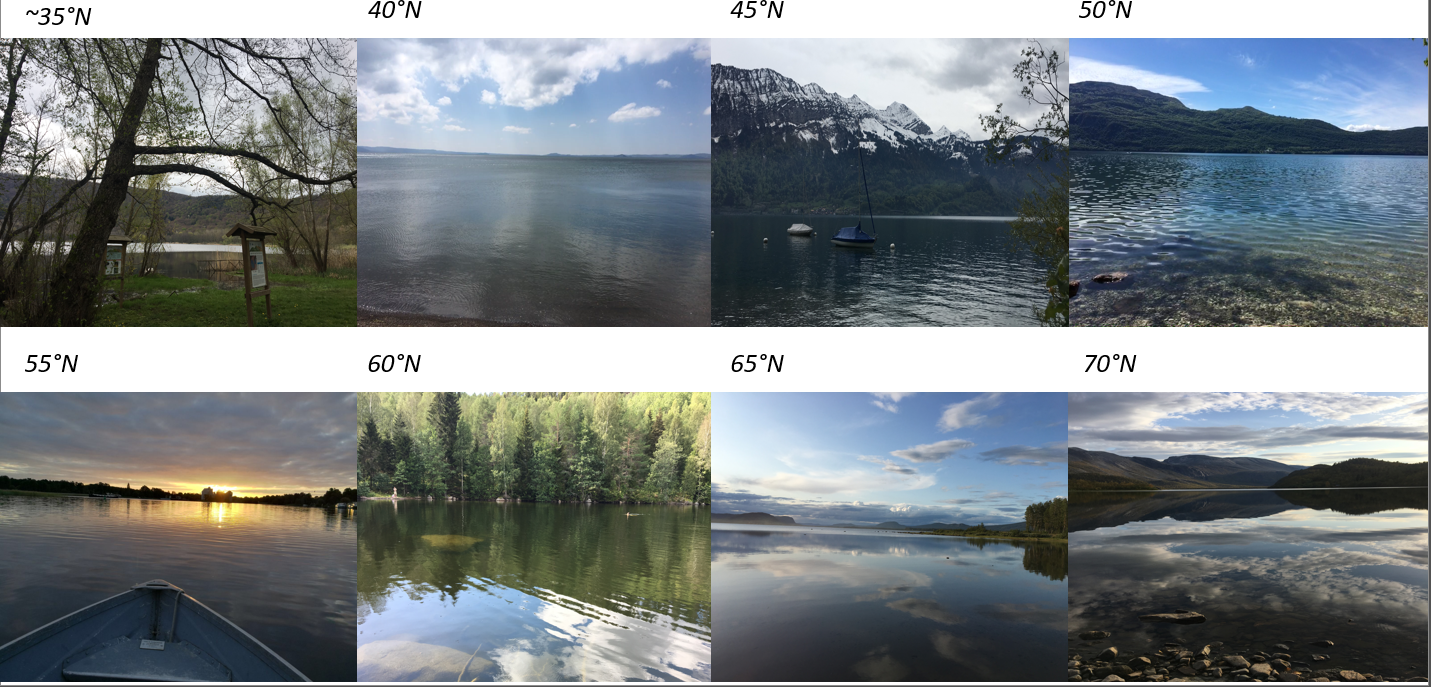

**Figure S18: Examples of lakes and their surrounding landscape along the studied latitudinal gradient.** At lower latitudes many lakes resulted from the accumulation of water in a depression formed by volcanic activities, as shown for Lago Monticchio Grande (~35°N) and Lago Albano (40°N). Lakes in Croatia resulted from sedimentation of calcium carbonate that form barriers and caused the formation of lakes (e.g. Lake Kozjak in Plitvice National Park). Other lakes resulted from the accumulation of water near mountains or higher terrain. On the southern side of the Alps, the catchment of those lakes primarily consisted of high elevation mountains with steep slopes and few trees, as illustrated by Grundlsee (45°N). We sampled few lakes between 50°N and 55°N that were located within the Vosges Mountain, as illustrated by Lac de Longemer (50°N) and other lakes near wetlands. At higher latitudes, most lakes were formed due to the last ice age that brought large volumes of ice on land. When those glaciers melted water accumulated and formed a lake. In northern Germany, those lakes were mainly surrounded by deciduous forests and farmlands, as illustrated by Röblinsee (55°N). In southern Scandinavia, most lakes were surrounded by evergreen forest, mostly pines, as illustrated by Tväringen (60°N). Several lakes in Scandinavia were also located near mountains and were similar to those found around the Alps. Lakes in the northern part of the gradient were remarkably shallow (few metres) for their size, as illustrated by Storavan (65°N). Further north, trees were largely inexistent, and lakes were surrounded by grass and shrubs on rocks, as illustrated by Nipivannet (70°N). All pictures were taken on the sampling day but not at the time of sampling, therefore the cloud cover and sunlight do not necessarily represent the conditions at the time of sampling.

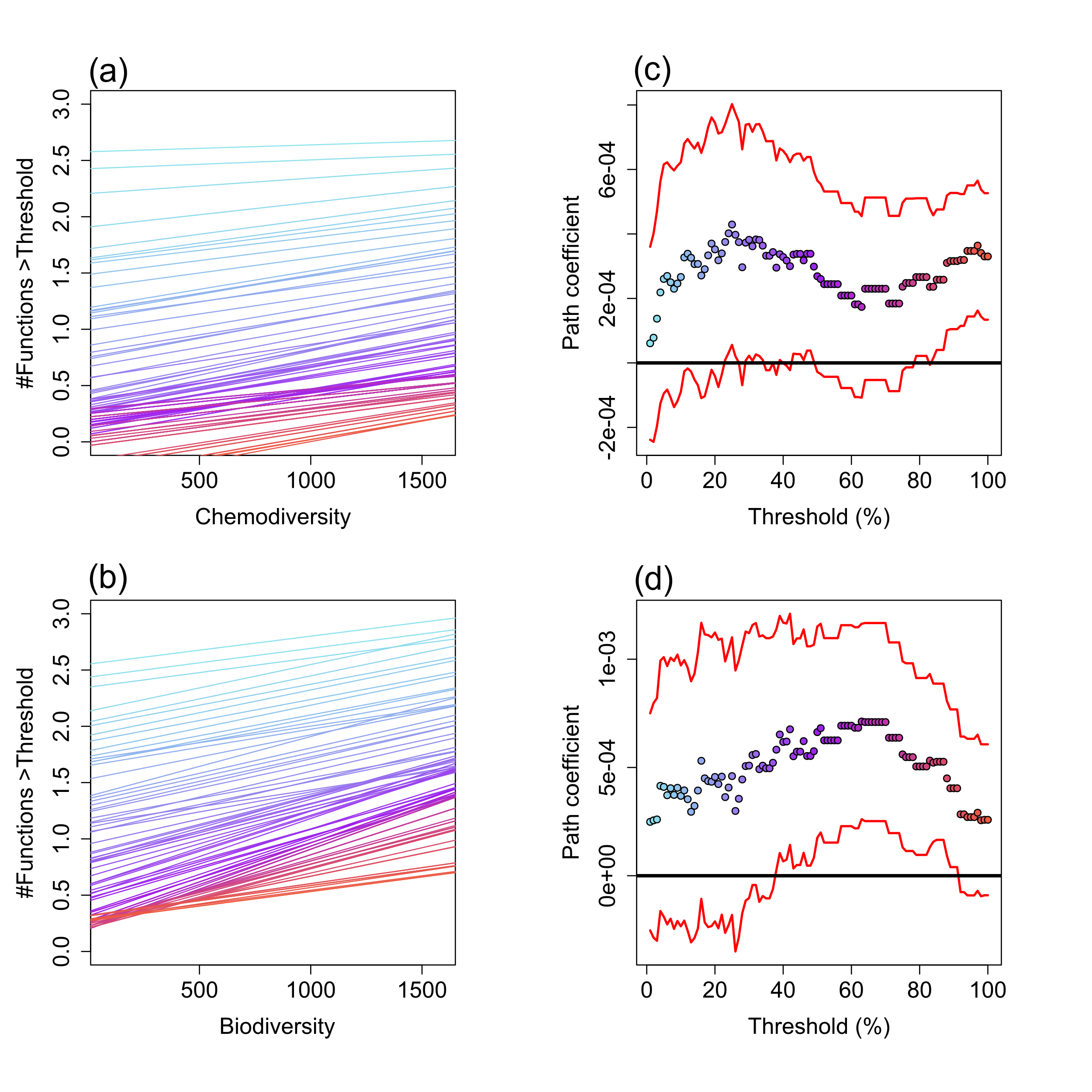

**Figure S19: Multifunctionality is supported more by chemodiversity than biodiversity when calculated using the threshold approach.** Lines are the estimated number of functions in the field survey exceeding a given threshold with increasing **(a)** chemodiversity or **(b)** biodiversity. We increased the threshold in 1% increments from 1% (light blue) to 99% (dark red) of the maximal values for each of the five measured functions. Colours correspond to points along the x-axis shown in **(c)** and **(d)**, which show mean (points) ± 95% confidence interval (lines) for slopes estimated between the number of functions above a given threshold and **(c)** chemodiversity or **(d)** biodiversity. In other words, the y-axis values of the points in **(c)** and **(d)** correspond to the slopes of the lines in **(a)** and (**b).** N=82. **Table S1: Characterization of the DOM used in the laboratory experiment**. For each source, we reported the number of assigned molecular formulae (#) along with the mean (standard deviation, SD) for their intensity-weighted molecular mass (m/z) and elemental ratios. We also calculated the proportion of formulae in each source that could be classified into different putative compound classes.

| **DOM source** | **#** | **m/z**  **(SD)** | **H:C**  **(SD)** | **O:C**  **(SD)** | **Aromatic** | **Unsatur-ated** | **Highly saturated** | **Saturated** |
| --- | --- | --- | --- | --- | --- | --- | --- | --- |
| Boreal lake | 3780 | 392.3  (129.1) | 1.12  (0.28) | 0.44  (0.16) | 9.8 | 12.7 | 77.5 | 0.1 |
| Arctic River | 2239 | 378.7  (126.1) | 1.23  (0.24) | 0.43  (0.15) | 18.7 | 9.7 | 71.4 | 0.2 |
| Temperate marsh | 3822 | 427.5  (144.0) | 1.01  (0.29) | 0.48  (0.16) | 30.1 | 4.7 | 64.9 | 0.3 |

**Table S2: Environmental variables were associated with chemodiversity.** We used a Spearman rank correlation to test the association between variables. N=101.

|  | **Chemodiversity** | |
| --- | --- | --- |
| **Variable** | **ρ** | **p** |
| NPP | -0.56 | <0.001 |
| Mean annual temperature | -0.46 | <0.001 |
| Mean annual precipitation | -0.42 | <0.001 |
| Water temperature | 0.23 | 0.015 |
| Water colour | 0.41 | <0.001 |
| pH | -0.18 | 0.052 |
| Lake area | -0.05 | 0.619 |
| Total dissolved nitrogen concentration | -0.14 | 0.160 |
| Chlorophyll a concentration | -0.13 | 0.230 |

**Table S3: Variable selection to predict chemodiversity.** At each step, we removed the least supported variable, that is, most decreased the Akaike information criterion (AIC) when removed from the best supported model. We repeated this process until removing a variable no longer reduced AIC by <2 units compared to the best supported model. Bolded parameters correspond to the variable that was dropped at each stepwise iteration.

| **Model** | **Number of parameters left after iteration** | **ΔΑΙC** |
| --- | --- | --- |
| Chemodiversity predictors: Net primary productivity + Precipitation + Mean annual temperature + Chlorophyll *a* + pH + Lake area + Water temperature + Latitude + Total dissolved nitrogen + Mean depth | | |
| Full model | 11 |  |
| **-Average depth** | 10 | -1.8 |
| **-Total dissolved nitrogen** | 9 | -2.0 |
| **-Latitude** | 8 | -1.9 |
| **- Water temperature** | 7 | -0.5 |
| **- Lake Area** | 6 | -0.8 |
| **- Mean annual temperature** | 5 | -1.6 |
| **- pH** | 4 | 0 |
| Final chemodiversity predictors: Net primary productivity + Precipitation + Chlorophyll *a* | | |

**Table S4: Model coefficients for the three ecosystem functions associated with chemodiversity**. Values are mean estimated effects (standard deviation) included in the best supported models, with bold indicating coefficients that differed statistically from 0. The coefficient of determination (R^2^) is provided for each model.

| **Coefficient** | **CH_4_** **emissions** | **CO_2_ emissions** | **Bacterial protein production** |
| --- | --- | --- | --- |
| Intercept | **0.53 (0.05)** | **90.15 (26.10)** | **-1.56 (0.04)** |
| Chemodiversity | **0.18 (0.07)** | **-71.02 (34.47)** | **0.20 (0.05)** |
| Mean annual temperature | **0.20 (0.07)** | -28.83 (30.76) | NA |
| Water colour | NA | **156.80 (86.52)** | **0.32 (0.06)** |
| Chlorophyll a | **0.12 (0.05)** | **-90.87 (30.61)** | **0.11 (0.05)** |
| Lake area | **-0.17 (0.05)** | NA | NA |
| R^2^ | 0.46 | 0.22 | 0.45 |
